# Evolutionary loss of inflammasomes in carnivores to facilitate carriage of zoonotic infections

**DOI:** 10.1101/2020.12.07.398529

**Authors:** Z. Digby, J. Rooney, J.P. Boyle, B. Bibo-Verdugo, R. Pickering, S.J. Webster, T.P. Monie, L.J. Hopkins, N. Kayagaki, G.S. Salvesen, S. Warming, L. Weinert, C.E. Bryant

**Affiliations:** Department of Veterinary Medicine, The University of Cambridge, Cambridge, CB30ES, UK; Sanford Burnham Prebys Medical Discovery Institute, 10901 North Torrey Pines, La Jolla, CA, USA 92037; Department of Physiological Chemistry, Genentech Inc., South San Francisco, California 94080, USA; Department of Molecular Biology, Genentech Inc., South San Francisco, California 94080, USA; University of Cambridge School of Clinical Medicine, Box 111, Cambridge Biomedical Campus, Cambridge, CB2 0SP, UK

## Abstract

Zoonotic infections, such as COVID-19, reside in animal hosts before jumping species to humans. The Carnivora, like mink, carry many zoonoses yet how diversity in host immune genes across species impact upon pathogen carriage are poorly understood. Here we describe a progressive evolutionary downregulation of pathogen sensing inflammasome pathways in Carnivora. This includes the loss of nucleotide-oligomerisation domain leucine rich repeat receptors (NLRs), acquisition of a unique caspase−1/−4 effector fusion protein that processes gasdermin D pore formation without inducing lytic cell death and the formation of an NLRP3-caspase-8 containing inflammasome that inefficiently processes interleukin-1β (IL-1β). Inflammasomes regulate gut immunity, but the carnivorous diet is antimicrobial suggesting a tolerance to the loss of these immune pathways. The consequences of systemic inflammasome downregulation, however, can reduce the host sensing of specific pathogens such that they can reside undetected in the Carnivora.

## Introduction

Viral and bacterial zoonotic pathogens, such as COVID-19 and *Salmonella* species, infect asymptomatic or symptomatic animal hosts which facilitate the transmission to humans. Pathogen genomics has yielded important discoveries about the diversity of different microorganisms in the context of disease (Weinert et al., 2015). Comparative biology of animal immune systems and their links to infection susceptibility are less well understood. This is partly due to a lack of tools, for example antibodies or other resources that make immune studies tractable, but the use of CRISPR/Cas9 gene editing is a universal technique that can be applied to cells from many animals. Approximately 49% of all carnivore species (for example mink and dogs), the highest proportion of any mammal order including bats, carry one or more unique zoonotic pathogens (Han et al., 2016). Whether this is because Carnivora are a large group of animals harbouring many pathogens so carry proportionally more zoonoses (Mollentze and Streicker, 2020) or due to other factors such as differences in the immune system remains to be determined.

Inflammasomes are of central importance in host protection against viral and bacterial diseases driving inflammation to control infections in humans and mice (Broz and Dixit, 2016). Canonical inflammasomes are a multi-protein complex composed of a pathogen recognition receptor, such as a nucleotide-oligomerisation domain leucine rich repeat receptors (NLRs; NLRP1, NLRP3 and NLRC4), Pyrin or absent-in-melanoma 2 receptor (AIM2), an adaptor (apoptosis associated speck like protein, ASC) and an effector protein (caspase-1; CASP1). The role of this pro-inflammatory protein complex is to process the immature cytokines pro-interleukin-1β (IL-1β) and pro-IL-18 into their mature, more active forms and to cleave the lytic pyroptotic cell death effector gasdermin D to its pore forming N-terminal fragment (Broz and Dixit, 2016). Non-canonical inflammasomes can also be formed by cytosolic delivery of the bacterial toxin lipopolysaccharide (LPS) which leads to the activation of caspase-11 in mice or caspase-4 and −5 in humans to cleave gasdermin D, which in turn leads to the activation of NLRP3 (Broz and Dixit, 2016; Broz et al., 2020; Lieberman et al., 2019). There is wide species diversity in AIM-2 like receptors, but there is substantial functional redundancy which acts as a compensatory mechanism (Brunette et al., 2012).

Here by comparing the distribution and evolution of inflammasome and cell death genes across the order Carnivora we find a profound compromise in inflammasome functionality, caspase-dependent lytic cell death pathways and a critical loss of NLR genes. A caspase-1-caspase-4 fusion protein found in all Carnivora despite being functionally capable of processing substrates *in vitro* and being active in mouse cells is inactive in cells from a model carnivore (dog). Caspase-8 is conserved and processes some pro-IL-1β, but without inducing lytic cell death and is predominantly required for upregulating expression of this protein. This compromised inflammasome activity coupled to the absence of the necroptotic effector Mixed Lineage Kinase Domain Like Pseudokinase (MLKL) (Dondelinger et al., 2016) should mean that the order Carnivora are immunologically challenged particularly in gut mucosal immunity, but ecology studies suggest that a high protein diet, such as that consumed by carnivores, is antimicrobial. This may explain why these innate immune pathways have been lost in the Carnivora, but the consequences for the carriage of zoonotic pathogens, particularly in organs other than the gut, may be very detrimental.

## Results

### The Carnivora caspase−1/−4 fusion has limited activity within cells compared to the recombinant protein and when an equivalent mouse caspase−1/−11 is expressed within mouse cells

Carnivora, such as dogs, cats and mink, through their close proximity with humans, can be susceptible to human pathogens. There are, however, marked differences in Carnivora inflammasome effector caspases compared to humans and also to mice. A unique caspase−1/−4 fusion protein is present in all cats and dogs (Eckhart et al., 2008). This protein has the equivalent of the CARD1 domain of caspase-1, which should render it unable to respond to cytosolic LPS, and the catalytic domain is most closely related to the catalytic domain of mouse caspase-11 (caspase 4 or 5 in humans), being only distantly related to caspase-1 in terms of sequence identity (Fig S1). This caspase−1/−4 fusion is conserved across all Carnivora (Fig 1A). The absence of individual caspase-1 and caspase−4/−5/−11 genes in Carnivora suggests there will be differences in how inflammasomes function in species of this order. The structural differences in the caspase−1/−4 fusion (Fig S1) suggests it should be able to process gasdermin D in response to canonical inflammasome stimulation, but have a limited/no capacity to process IL-1β and IL-18. To characterise inflammasome functionality in Carnivora we first took immortalised bone marrow derived macrophages from wild type (WT) mice and compared them to the DH82 dog macrophage-like cell line as a Carnivora model. Cells were infected with *Salmonella enterica* serovar Typhimurium (*S*. Typhimurium) which activates NLRC4 and NLRP3 canonical inflammasome formation (Man et al., 2014). IL-1β was produced by the WT mouse cells, as expected, and also, surprisingly, the dog cells which showed clear, but inefficient, processing of pro-IL-1β (Fig 1B). Cells from WT mice, caspase−1/−11 double knock out mice and the DH82 dog cells infected with *S*. Typhimurium showed that the majority of cells from the WT mice, as expected, released LDH so lysed 2 hours post infection and cells from the caspase−1/−11 double knock out do not lyse until 24 hours post infection. The dog macrophage cells, surprisingly, did not lyse until 24 hours post infection (Fig 1B). Experiments in primary monocyte derived macrophages isolated from dog peripheral blood mononuclear cells (PBMCs) infected with *S*. Typhimurium also induced IL-1β production with delayed lytic cell death (Fig 1C). This is unexpected given the caspase−1/−4 fusion was expected to process gasdermin D to lyse cells, but not cleave IL-1β in response to canonical inflammasome activity (Kayagaki et al., 2011).

**Figure 1.**
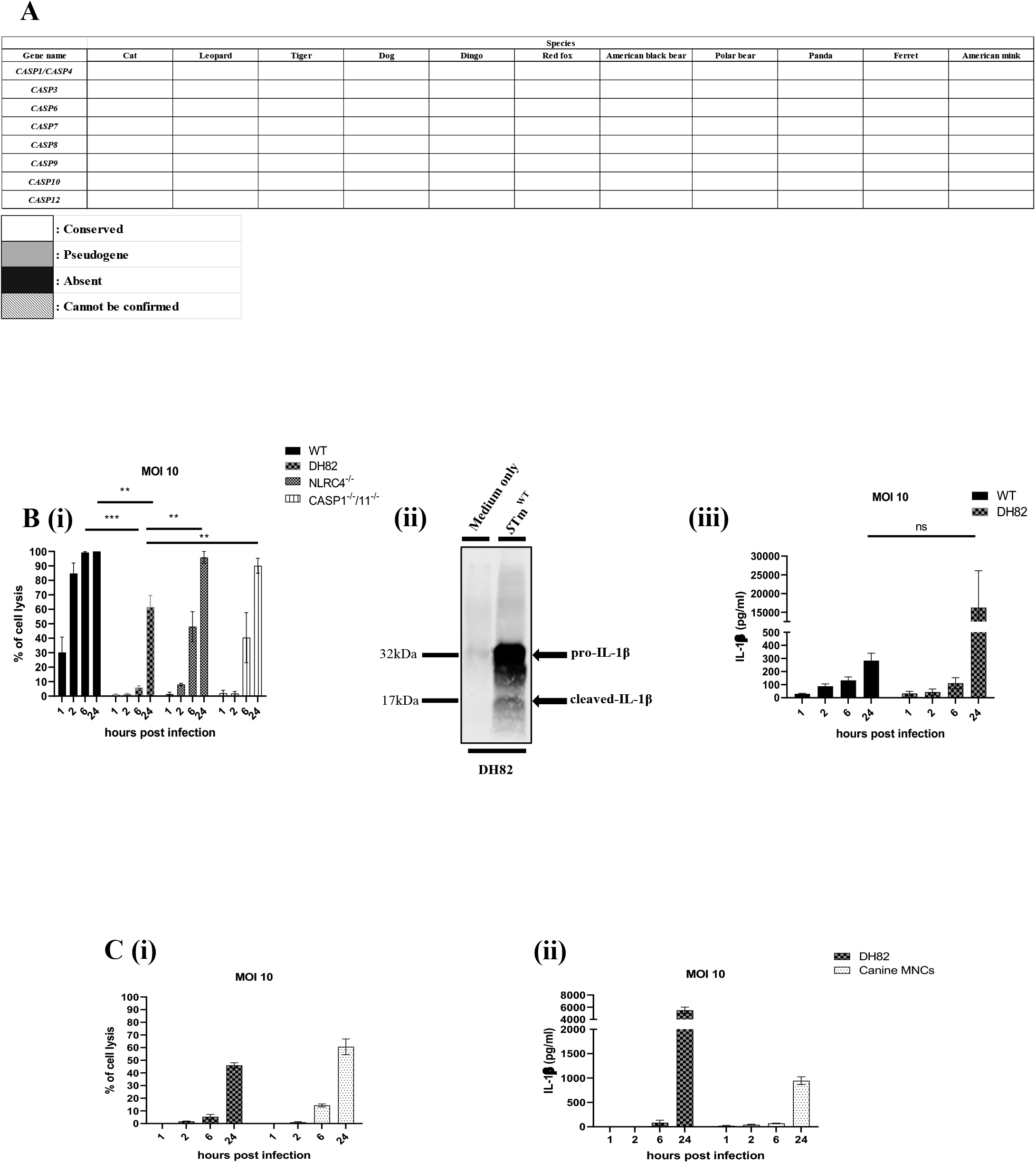
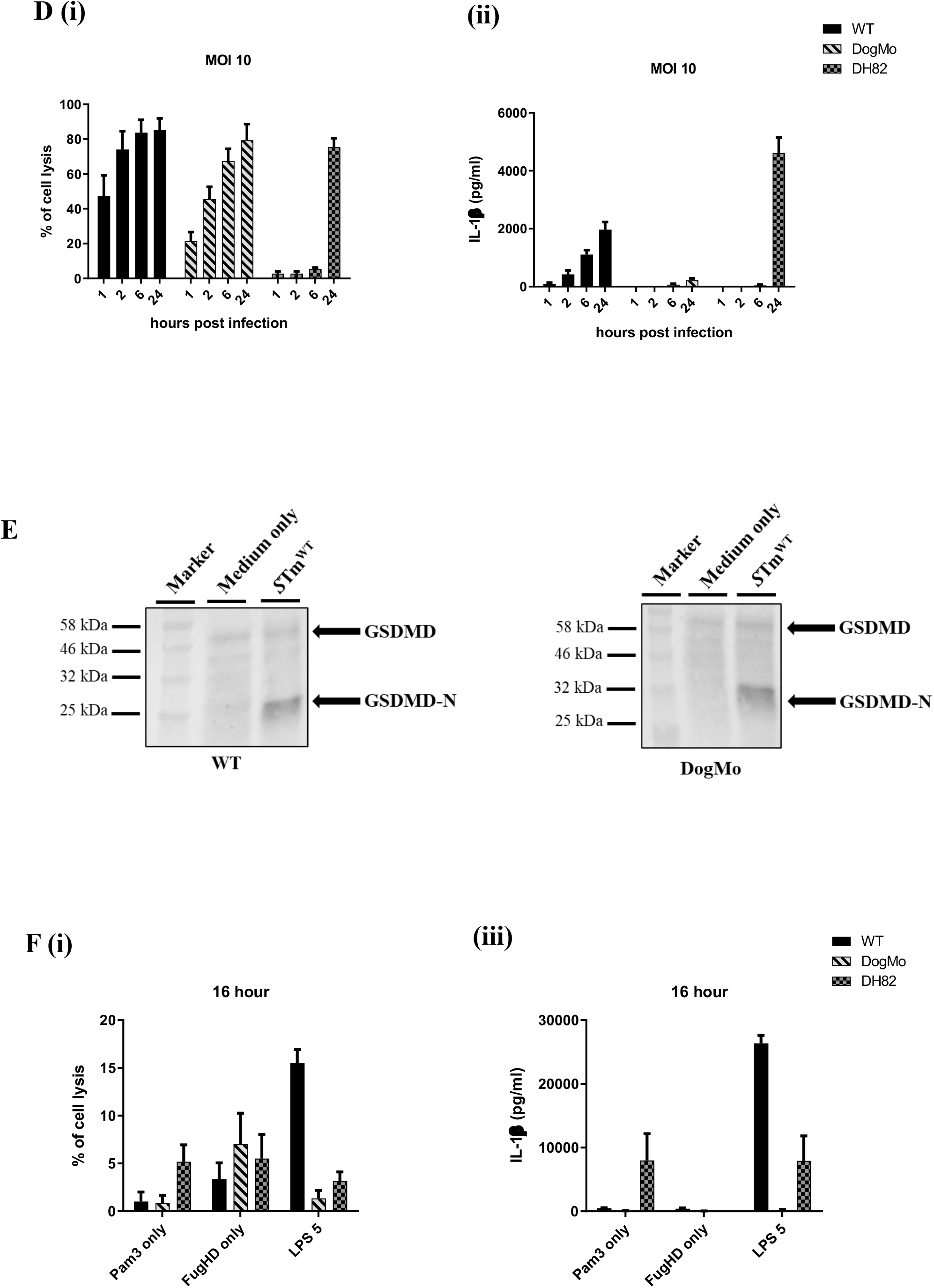
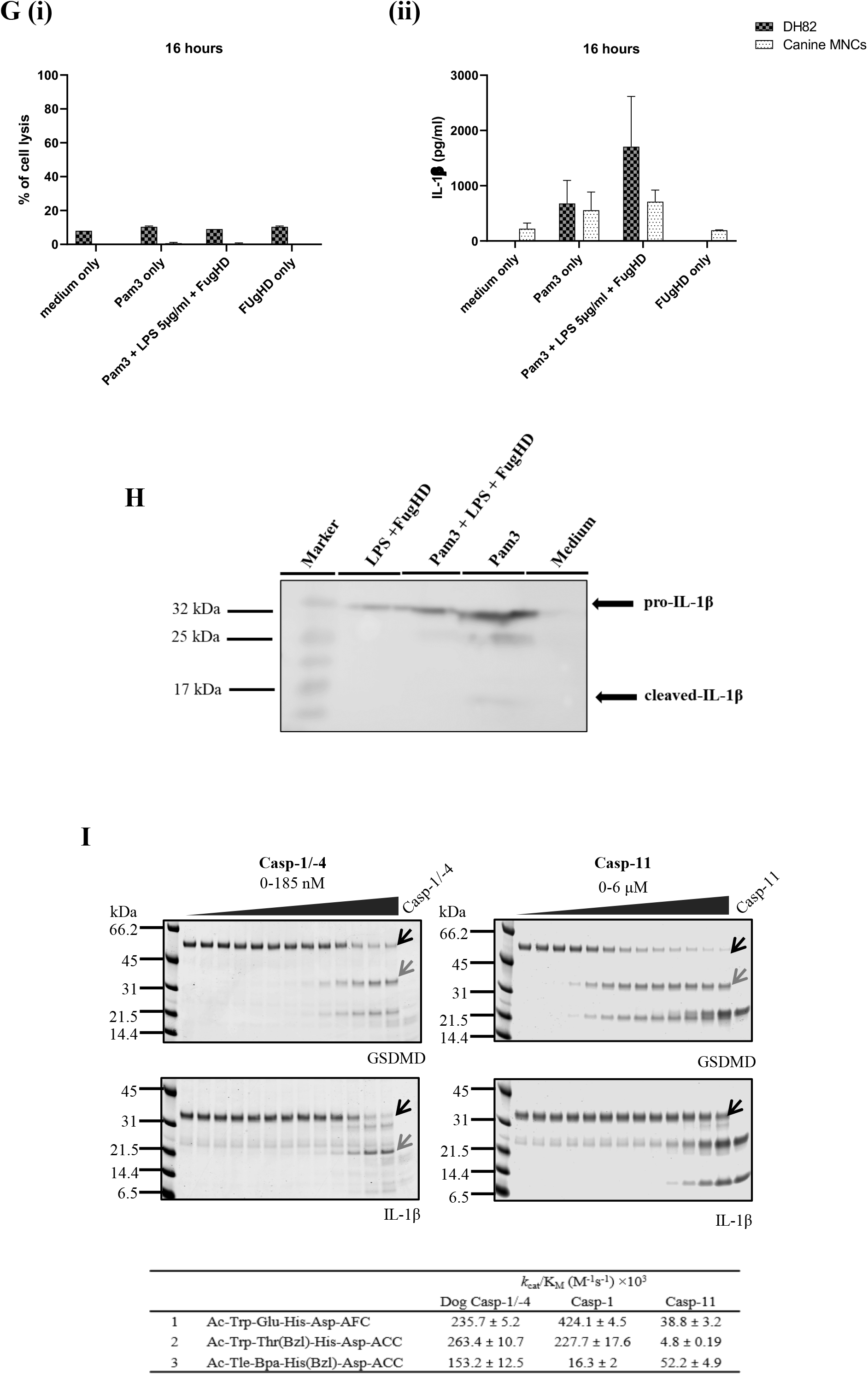
The Carnivora caspase−1/−4 fusion has limited activity within cells compared to the recombinant protein and when an equivalent mouse caspase−1/−11 is expressed within mouse cells. (A) Representative table showing the evolutionary conservation of the inflammatory hybrid caspase−1/−4 and apoptotic caspases in species belonging to order Carnivora. (Bi) Immortalised murine WT, NLRC4^*−/−*^, CASP1^*−/−*^/11^*−/−*^, canine DH82, (C) canine primary canine MNCs and immortalised DH82, (E) primary murine WT, DogMo and canine immortalised DH82 cells were infected with *S*. Typhimurium MOI of 10. (G) Murine primary WT, DogMo, canine immortalised DH82 cells and (H) canine primary MNCs and immortalised DH82 cells were primed with Pam3CSK4 (10 μg/ml for 4 hours) then transfected with LPS at 5 μg/ml final concentrations using Fugene HD. Percentage of cell lysis (Bi, Ci, Ei, Gi, Hi) and the amount of IL-1β released (Cii, Eii, Gii, Hii) into the supernatant at 1, 2, 6 and 24 hour post infections were determined via the measurement of relative LDH release and ELISA detection method, respectively. (Bii) Full protein extraction of DH82 cell lysates infected with *S*. Typhimurium MOI of 1 for 24 hours and (I) Pam3CSK4 (10 μg/ml for 4 hours) primed LPS transfected (5 μg/ml for 16 hours) were blotted against canine specific IL-1β (Bii) alongside non-infected controls (shown as medium only). (D) Upper: *in vitro* protein cleavage. Recombinant mouse gasdermin D and pro-IL-1β were submitted to cleavage with a dilution series of dog caspase−1/−4 or mouse caspase-11, incubated for 30 minutes and followed with SDS-PAGE analysis of cleavage products. Black arrows indicate bands corresponding to intact substrate and grey arrows correspond to the biologically active form of the protein originated by caspase cleavage. Lower: Catalytic efficiency represented as kcat/KM (M-1s-1) values for peptidyl fluorogenic substrates (1) general inflammatory caspase substrate, (2) caspase-1 selective substrate, or (3) caspase-11 selective substrate. Average of three determinations ± SD. (F) Murine primary WT and DogMo cells were infected with *S*. Typhimurium usingMOI of 10 for 6 hours. GSDMD cleavage was determined by western blotting of cell lysates using anti-mouse GSDMD antibody. Data shown is pooled from three (B), two (E, G) and a single (cells pooled from 4-6 dogs (Table S1) C, F, H) independent experiment(s). Error bars represent the standard error of means (SEM) of triplicate wells

Caspase 1 and caspase 11 are so close to each other in the mouse genome (Kayagaki et al., 2011) that it is theoretically possible to delete the catalytic domain of caspase 1 and the CARD domains of caspase 11 to make a mouse fusion protein equivalent to the Carnivora caspase−1/−4 fusion. To determine the biological functions of the caspase−1/−4 fusion we therefore used a novel approach of CRISPR/Cas9 gene deletion (S2) to generate a mouse with a fusion protein of the caspase-1 CARD1 and the caspase-11 catalytic domain (DogMo; (Fig S3). Bone marrow derived macrophages from DogMo infected with *S*. Typhimurium, as expected, showed canonical inflammasome driven cell lysis and gasdermin D processing, but no IL-1β production (Fig 1D, E). DogMo cells also showed no cell lysis or IL-1β production in response to non-canonical inflammasome activation induced by cytosolic LPS (Fig 1F). DH82 dog cells and primary dog mononuclear cells, in contrast, showed no cell lysis and limited IL-1β production in response to cytosolic LPS (Fig 1F, G, S4). The IL-1β produced by the DH82 cells was the same whether the cells were stimulated with the TLR2 ligand Pam3CSK4 (as a control) or after transfection of LPS into the cytosol after priming with Pam3CSK4 suggesting this cytokine was induced by priming rather than non-canonical inflammasome activation (Fig 1F, G, H). To determine whether the enzymatic properties of the caspase−1/−4 hybrid could account for the IL-1β processing in the absence of lytic cell death by inflammasomes in dog cells, we expressed the catalytic domain of this protein to test its ability to process substrates *in vitro* (Fig 1I). Caspase−1/−4 processed both gasdermin D and IL-1β to their biologically active forms *in vitro*. Caspase-11, as expected, cleaved gasdermin D but not IL-1β. We also used synthetic peptidyl substrates optimized, based on specificity screens, to improve selectivity for caspase-1 or caspase-11 (Ramirez et al., 2018). Dog caspase−1/−4 cleaved these synthetic substrates at superior rates compared to caspase-11 and cleaved the caspase-1 optimum substrate at rates comparable to caspase-1 (Fig 1I). These data reveal that the substrate specificity of dog caspase−1/−4 resembles that of caspase-1 more closely than caspase-11 *in vitro.* This suggests that the defective canonical and non canonical inflammasome responses in dog cells are not caused by an intrinsic loss of enzymatic activity of the caspase−1/−4 fusion protein, but most likely because of an alternative regulatory mechanism.

### The inflammasome gene repertoire and functionality in the Carnivora is severely compromised

An alternative explanation for our data is that species-specific differences in NLRs might account for the weak activation of canonical inflammasomes we see in dog cells. AIM2, for example, is absent in many species including Carnivora (Brunette et al., 2012). Analysis of the repertoire of NLR genes across the Carnivora identified that either NLR family of apoptosis inhibitory proteins (NAIP) and/or NLRC4 (bacterial sensors) are predominantly missing or are pseudogenes in the Canidae whereas the Felidae lack another bacterial sensor NLRP1 (Fig 2A). The loss of these NLRs occurred early in the evolutionary trees of both the Canidae and Felidae (Fig 2B). This diversity in NLRs suggests that the lack of NAIP/NLRC4 in dogs would, at least partially, explain the altered inflammasome responses to *S*. Typhimurium, which activates both NLRC4 and NLRP3, we saw when comparing mouse and dog macrophages. NLRP3, a non-specific sensor of cellular insults (Swanson et al., 2019), in contrast, is conserved across all Carnivora. We next stimulated WT mouse, caspase−1/−11 double knock out mouse or dog DH82 macrophages with increasing concentrations of the canonical NLRP3 activator nigericin. We saw cell lysis in WT mouse BMMs, as expected, but no cell lysis in dog cells until we used very high concentrations of nigericin when cell death was independent of both caspase-1 and −11 in mouse BMMs (Fig 2C). The failure of nigericin to induce inflammasome-induced cell lysis was not due to the inability of the caspase fusion protein to function per se because stimulation of the DogMo macrophages again showed cell lysis and gasdermin D cleavage, but minimal IL-1β production (Fig 2D, E).

**Figure 2.**
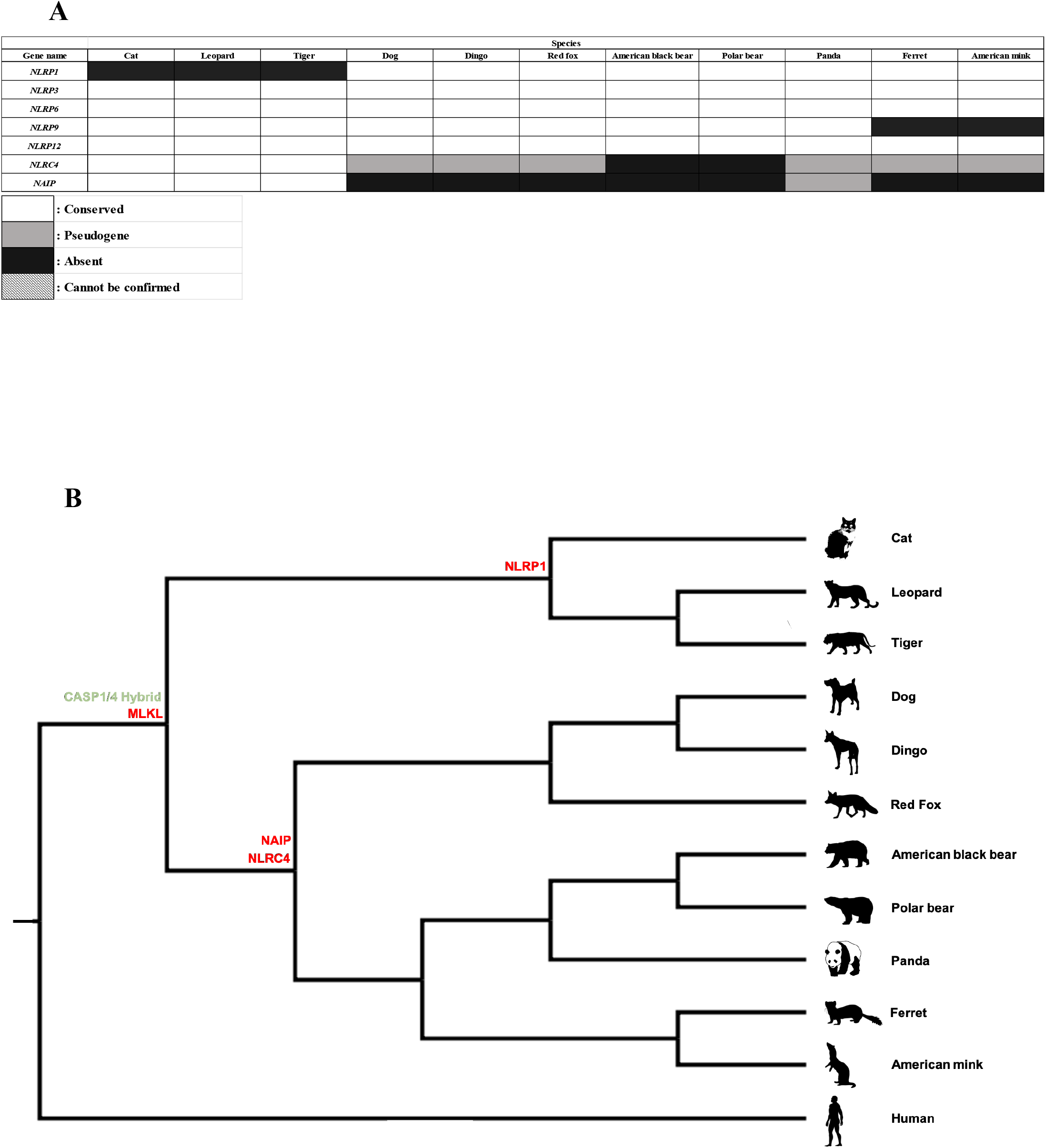
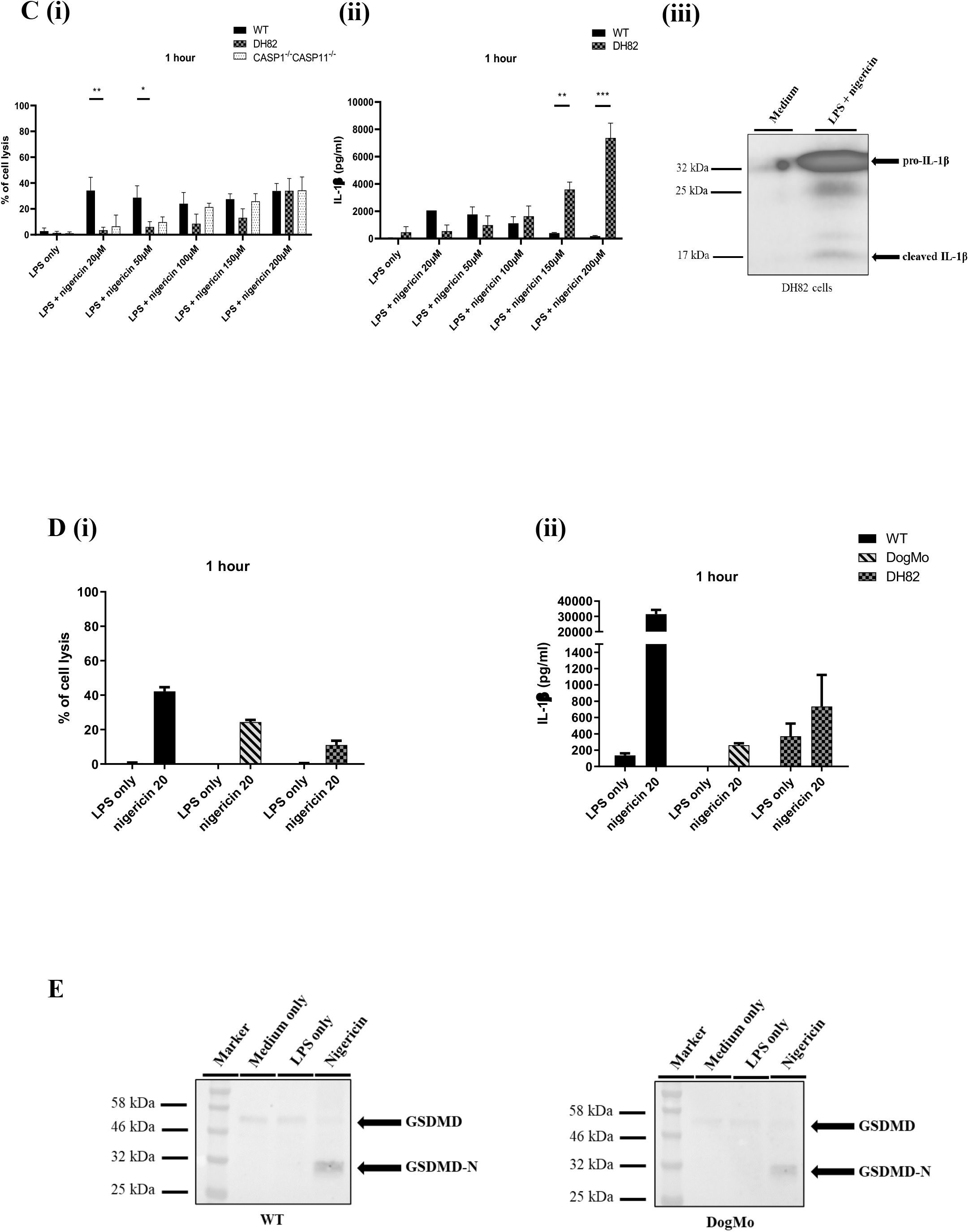
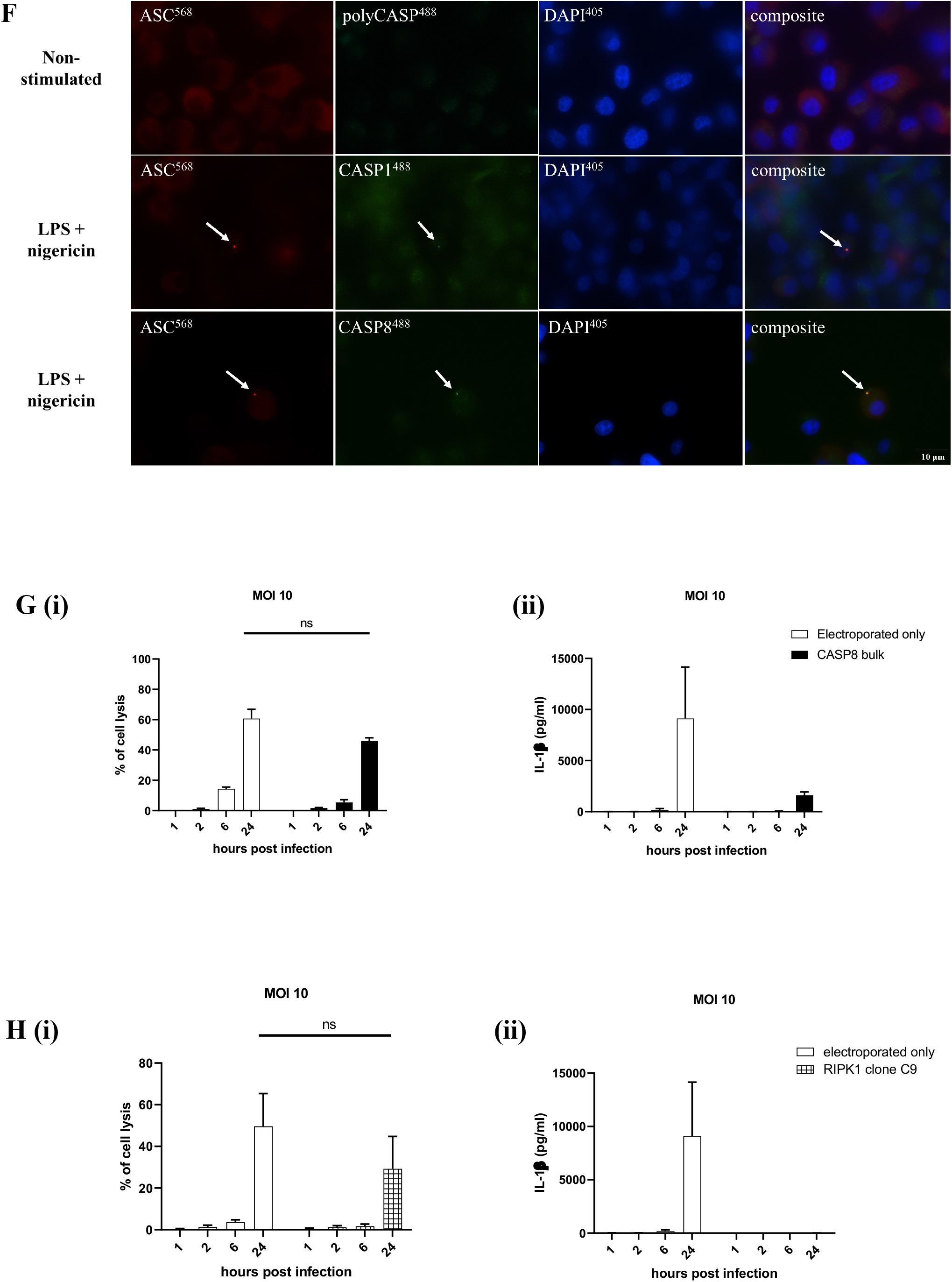
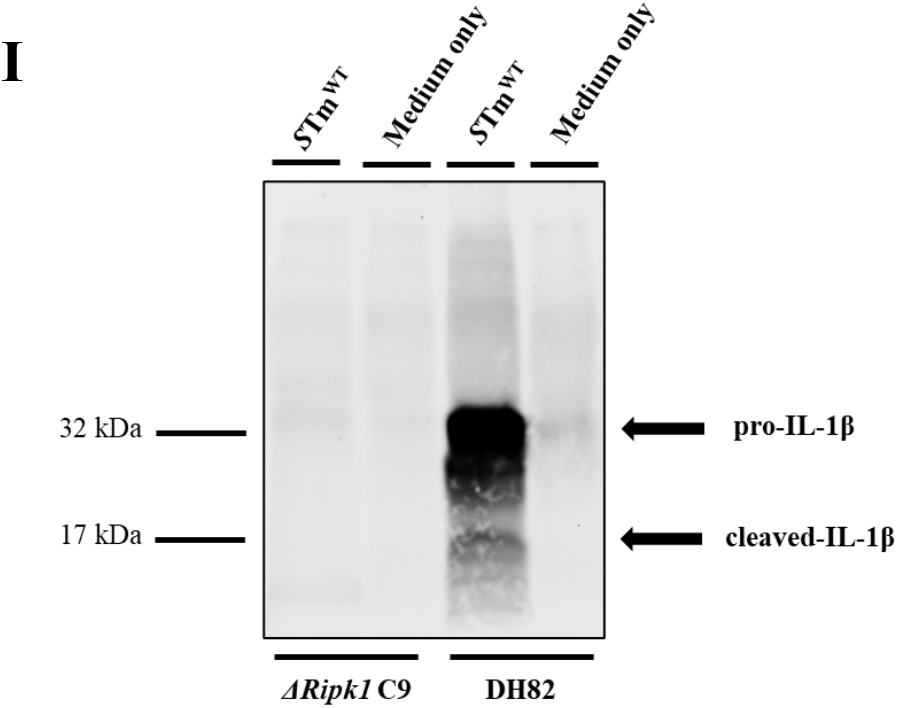
The inflammasome gene repertoire and functionality in the Carnivora is severely compromised. (A) Representative table showing the evolutionary conservation of pattern recognition receptors (PRRs) in species belonging to order Carnivora. (B) Species tree of the order Carnivora available on the Ensembl database showing gain and loss of genes. Gene loss is represented by gene names labelled in red, while gene gain (or substitution) is represented by gene names labelled in green. (C) Immortalised murine WT, CASP1^−/−^CASP11^−/−^, canine immortalised DH82, (D) primary murine WT, DogMo and immortalised canine DH82 cells were primed with LPS (200 ng/ml for 3 hours) then stimulated with nigericin (20-200 μM for 1 hour). Percentage of cell lysis (Ci, Di) was determined by measuring the relative amount of LDH release into the supernatant, while IL-1β release (Cii, Dii) was measured using murine- and canine-specific ELISA kits, respectively. (Ciii) Immortalised DH82, (E) primary WT and DogMo cells were primed with LPS (200 ng/ml for 3 hours), following stimulation with nigericin (200 μM and 20 μM for 1 hour, respectively). Total protein was precipitated from the supernatant and (Ciii) IL-1β and (E) Gasdermin D cleavage was investigated using canine- and murine specific IL-1β and Gasdermin D antibodies, respectively using western blotting. (F) DH82 cells were primed with LPS (200 ng/ml) and stimulated with nigericin (20 μM for 1 hour). Non-stimulated (NS) cells were left in cell culture medium. Live cells were stained for activated caspases (FLICA poly-caspase, FLICA caspase-1 or FLICA caspase-8 (green)). Following fixation cells were stained for cytoplasmic ASC (red) and nuclei using DAPI staining (blue). (G) Bulk-edited CASP8- and (H) RIPK1-deficient clones, WT and electroporated only WT DH82 cells were infected with *S*. Typhimurium at MOI of 10. Percentage of cell lysis (Gi, Hi) induced was determined by measuring the relative amount of LDH release into the supernatant, while IL-1β release (Gii, Hii) was measured using canine-specific ELISA kits. (I) WT and RIPK1-deficient DH82 cells were infected with *S*. Typhimurium at MOI of 1 for 24 hours. Full protein extraction of cell lysates were blotted against canine specific IL-1β. Non-infected controls (shown as medium only) were also included for each cell line. Data shown is pooled from three (C, F, G, I), two (D) and a single (E, I) independent experiment(s). Error bars represent the standard error of means (SEM) of triplicate wells

How then is NLRP3 processing IL-1β, but not driving cell lysis? Caspase-8 can be recruited to the inflammasome to process IL-1β and gasdermin D, particularly in the absence of caspase-1 (Lee et al., 2018; Man et al., 2013; Newton et al., 2019). In experiments where we visualised NLRP3 induction of ASC specks we saw recruitment of a caspase-8 fluorescent substrate to this macromolecular structure within dog cells (Fig 2F). To validate this observation, we used CRISPR/Cas9 to delete caspase-8 from dog macrophages (S5) and saw a loss of IL-1β production in response to *S*. Typhimurium (Fig 2G). This suggests canonical inflammasome formation in dog macrophages utilises caspase-8, rather than the caspase-1−4 fusion, to weakly process IL-1β (Fig 2I), but lytic cell death is not initiated. We also used a caspase-1 fluorescent substrate and visualised the caspase−1/−4 fusion recruited to the ASC speck (Fig 2F), but deletion of this gene fusion by CRISPR/Cas9 did not alter IL-1β production by DH82 cells following cellular infection with *S*. Typhimurium (S5). Caspase-8 regulates the Receptor-interacting serine/threonine-protein kinase 1 (RIPK1)/RIPK3 pathway that triggers necroptosis, but also regulates Toll-like receptor 4 (TLR4)-dependent Nuclear factor kappa-light-chain-enhancer of activated B cells (NF-κβ) driven transcription of genes such as pro-IL1β. Carnivora cells cannot undergo necroptosis as the effector protein Mixed Lineage Kinase Domain Like Pseudokinase (MLKL) is missing from their genomes (Dondelinger et al., 2016). When we used CRISPR/Cas9 to remove RIPK1 from dog cells IL-1β production was abolished in response to inflammasome stimulation (Fig 2H, I) because of a loss of pro-IL-1β transcription driven by TLR4 priming from the *Salmonella* LPS. Collectively our data suggest that what little inflammasome cytokine processing occurs in carnivores is driven by caspase-8.

### Inflammasome activity induces pore formation, but is uncoupled from cell lysis in the Carnivora

Our data suggest that in dogs and, presumably in other Carnivora, non-canonical inflammasome activation is absent and canonical inflammasome activation is limited to a little caspase-8-dependent processing of IL-1β without lytic cell death. In Carnivora there is a loss of, or modification of, genes important in lytic cell death pathways (necroptosis and pyroptosis Figure 3 A, E). Gasdermin D pores can form, however, without inducing lytic cell death (Evavold et al., 2018) so as there are no gasdermin D antibodies that cross react with the dog protein we measured propridium iodide (PI) uptake and used live cell imaging of DH82 cells to determine if pyroptotic cell death pathways are completely missing in dog cells. *S.* Typhimurium classically processes gasdermin D to induce rapid lytic cell death in mouse or human cells, yet infected dog cells took up PI, but only lysed 24 hours post infection presumably when the cell membrane can no longer contain the enormous intracellular bacterial load (Fig 3 B, C). In response to nigericin stimulation DH82 cells again took up PI suggesting gasdermin D pores are formed, but cells from these animals have a markedly reduced capacity for pro-inflammatory lytic cell death (Fig 3 B, D). The appearance of the caspase−1/−4 fusion and the loss of MLKL occurred early in the Carnivora evolutionary tree (Fig 2B). Gasdermin E, a protein that drives pyroptosis in response to caspase-3 activation (Wang et al., 2017), is conserved. The apoptotic caspases 3, 7, 8 and 9 are fully conserved across all Carnivora suggesting caspase-dependent cell death may be limited primarily to apoptosis pathways in these animals (Fig 3A). We do see some lytic cell death at very high doses of nigericin in dog cells (Fig 3C) which could be driven by, for example, gasdermin E, but this occurs under conditions of limited physiological relevance.

**Figure 3.**
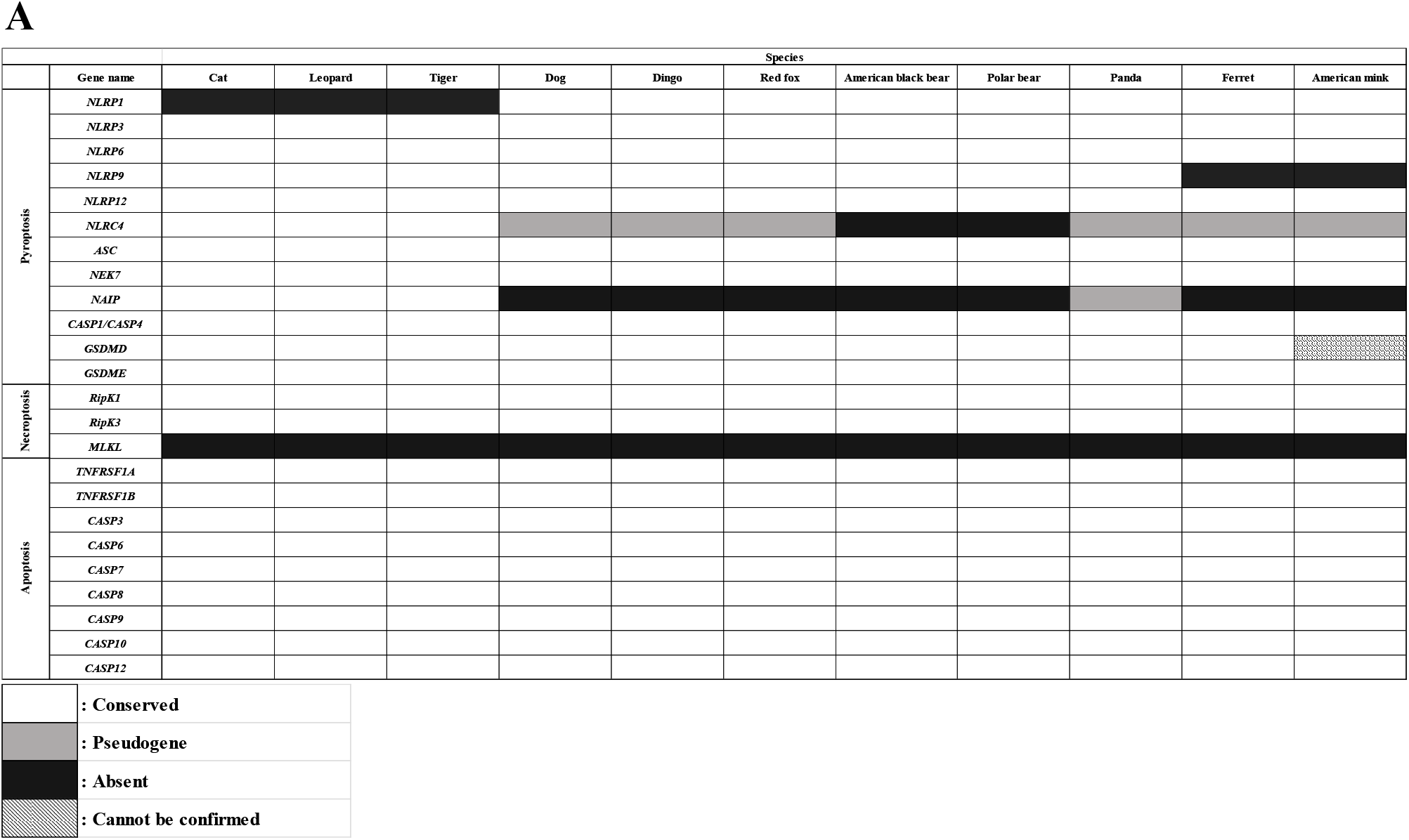
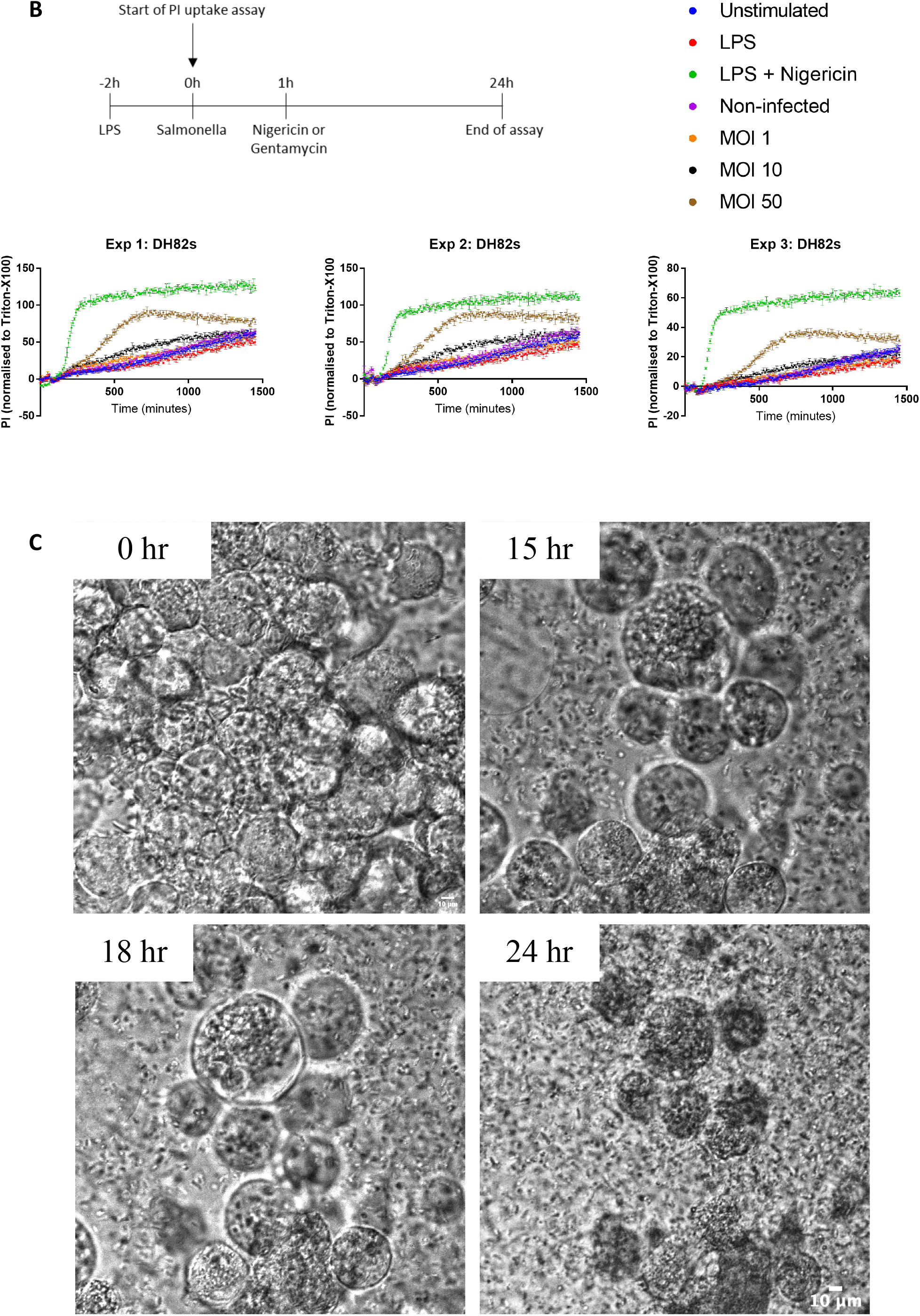
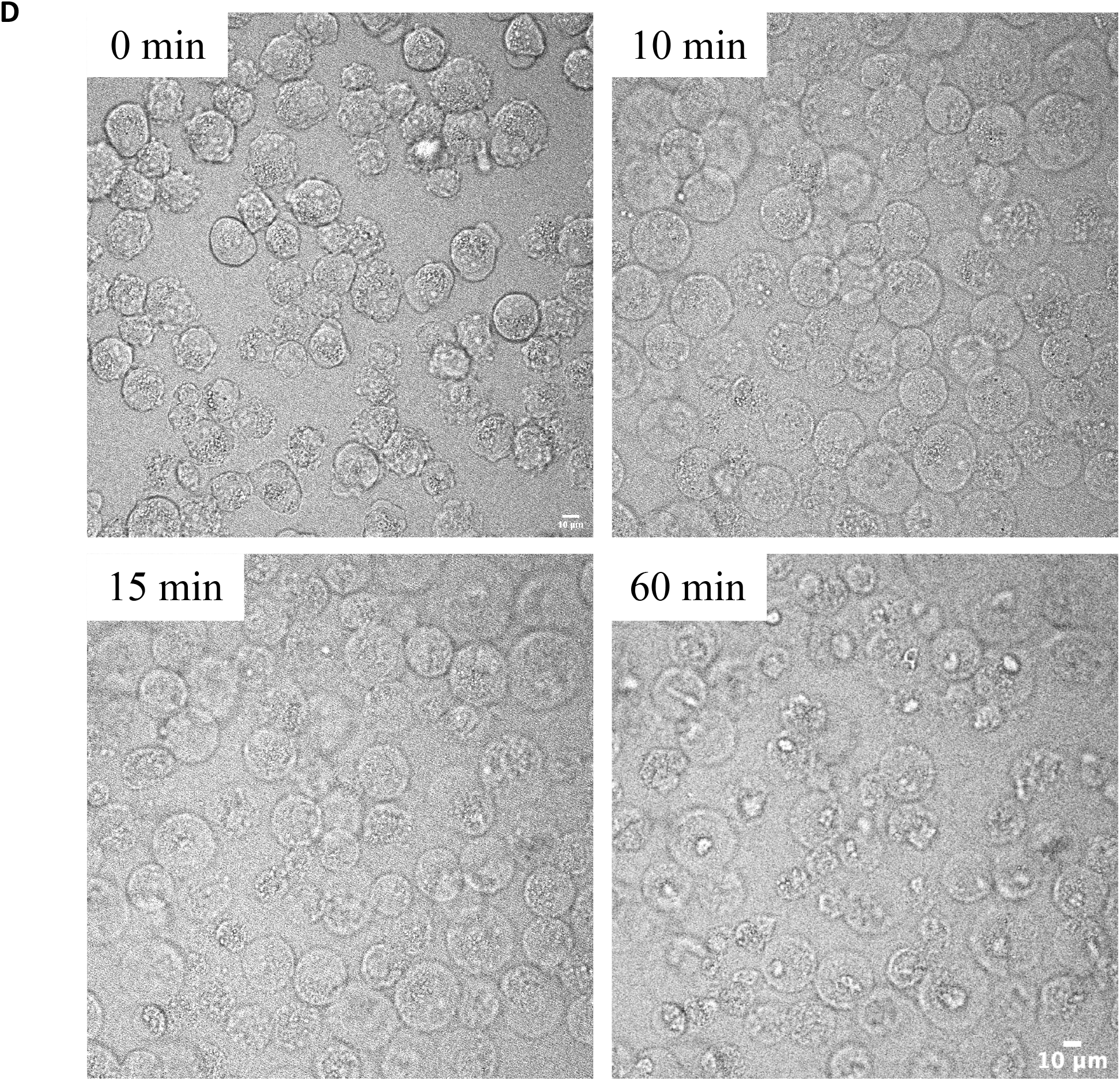
Inflammasome activity induces pore formation, but is uncoupled from cell lysis in the Carnivora. (A) Representative table showing the evolutionary conservation of main components of different lytic cell death pathways in species belonging to the Carnivora order. (B, C, D) DH82 cells were infected with WT *S*. Typhimurium MOI of 1-50 for 24 hours or primed with LPS (200 ng/ml) for 3 hours and stimulated with nigericin (200 μM) for 1 hour. Propridium iodide uptake was measured (B) or bright field images were taken throughout the incubation on a confocal microscope (every 5 minutes for the infection (C) and every minute for nigericin stimulation experiment (D)). Representative images of the different morphological stages observed are shown above from single experiments (movies M1 and M2 accompany this paper). Scale bar = 10μm

## Discussion

Here we show that key inflammasome lytic cell death pathways thought to be critical for gut health in mammals are either genetically and/or functionally missing from the Carnivora. PI uptake without cell lysis occurs in dog cells in response to inflammasome activation suggesting a dissociation of gasdermin D pore formation from lytic cell death similar to the phenotype seen if the NINJ1 protein is deleted from human or mouse cells (Kayagaki et al., 2020) yet all Carnivora have NINJ1. Our data suggest that inefficient inflammasome activation occurs in dog cells such that insufficient gasdermin D pores form to drive cell lysis. Inflammasome-driven lytic cell death is, therefore, lost in Canidae and this is particularly interesting because the Carnivora also lack the necroptotic effector MLKL such that two of the critical inflammatory cell death pathways that are thought to be essential for host protection against infection are absent. One of the key functions for inflammatory cell death is to protect the gut against infection (Crowley et al., 2020; Rauch et al., 2017; Schwarzer et al., 2020; Sellin et al., 2014; Tummers et al., 2020) yet canonical inflammasome, non-canonical inflammasome and necroptotic lytic cell death pathways in Carnivora are inactive. Emerging ecological evidence suggests that a high protein diet, such as that consumed by the Carnivora, is antimicrobial (Cotter et al., 2019) and we speculate that the evolutionary and functional reduction in inflammasome activation in the Carnivora may simply reflect the protection against infection conferred on these animals by their dietary habits.

Canonical inflammasomes traditionally couple a receptor, such as an NLR or AIM2, with ASC and the effector protein caspase-1 whereas non canonical inflammasomes are driven by caspase−4/−5/−11 to indirectly activate NLRP3 (Broz and Dixit, 2016). In the Carnivora caspase-1 and −4 are fused into a single caspase which has the CARD domain of caspase-1 but the catalytic site of caspase-4/−11 which should result in a protein that is unable to bind to LPS, efficiently processes gasdermin D, but cannot process pro-IL-1β. Macrophages from a mouse expressing an analogous caspase−1/−11 fusion fulfilled these predictions, although when expressed as a recombinant protein the dog caspase−1/−4 can process both gasdermin D and pro-IL-1β *in vitro*. In the dog macrophages, however, the caspase−1/−4 fusion is relative inactive producing no lytic cell death, limited gasdermin D pore formation and what little pro-IL-1β processing occurs is through caspase-8. This caspase can be recruited to canonical inflammasomes, particularly in the absence of caspase-1, to process pro-IL-1β and drive cell death (Man et al., 2013; Newton et al., 2019). Our data shows that the predominant role for caspase-8 in the Carnivora, however, is to drive pro-IL-1β expression. IL-1β is active in domestic members of the Carnivora, like dogs (Dinarello, 2018), so presumably inflammasome independent mechanisms, such as through proteinase-3, elastase, matrix metalloprotease-9 and granzyme A (Dinarello, 2018), process sufficient pro-IL-1β to provide immune protection in these animals.

There are potential consequences for the loss of the inflammasome and cell death pathways particularly in organ systems other than the gut. The presence of bacteria in other organs of the Carnivora might be expected to lead to either immune deficits and/or allow the pathogen to hide. Inflammasome-induced pyroptosis and subsequent lytic cell death is thought to be a critically important mechanism by which pathogens are controlled in the host (Martinon et al., 2009; Miao et al., 2010). The lack of pyroptotic cell lysis in the Carnivora should, therefore, severely compromise the susceptibility of these animals to infection, yet these mammals continue to thrive. The Canidae lack of NAIP/NLRC4 should affect the recognition of a number of bacterial pathogens including salmonellae, *Pseudomonas* and *Bacillus*. The Felidae lack of NLRP1, which is activated by protease cleavage and thus susceptible to processing by bacterial or viral proteases (Chui et al., 2019; Sandstrom et al., 2019), potentially compromises the immune status of these animals in response to pathogens. The Carnivora also have a minimal capacity to activate NLRP3.

The compromise in inflammasome functionality in the Carnivora does not seem to predispose them to infection, but another possible consequence of the lack of PRR recognition could be to allow pathogens to reside undetected facilitating the potential for zoonotic carriage. One reason for the high zoonotic carriage by bats has been suggested to be due to dampened activation of NLRP3 (Ahn et al., 2019). The muted inflammasome activation we see in Canidae might also facilitate zoonotic carriage. COVID-19, for example, is carried by mink (Oude Munnink et al., 2020) and although its genome is incomplete the caspase 1 locus is intact with, as expected, a predicted caspase−1/−4 fusion protein consistent with poor inflammasome functionality. The loss of these immune pathways in Carnivora may, therefore, have the unexpected consequence of permitting certain pathogens to evade host detection when in habitats other than the gut thus facilitating the carriage of zoonotic infections.

**Table 1.**
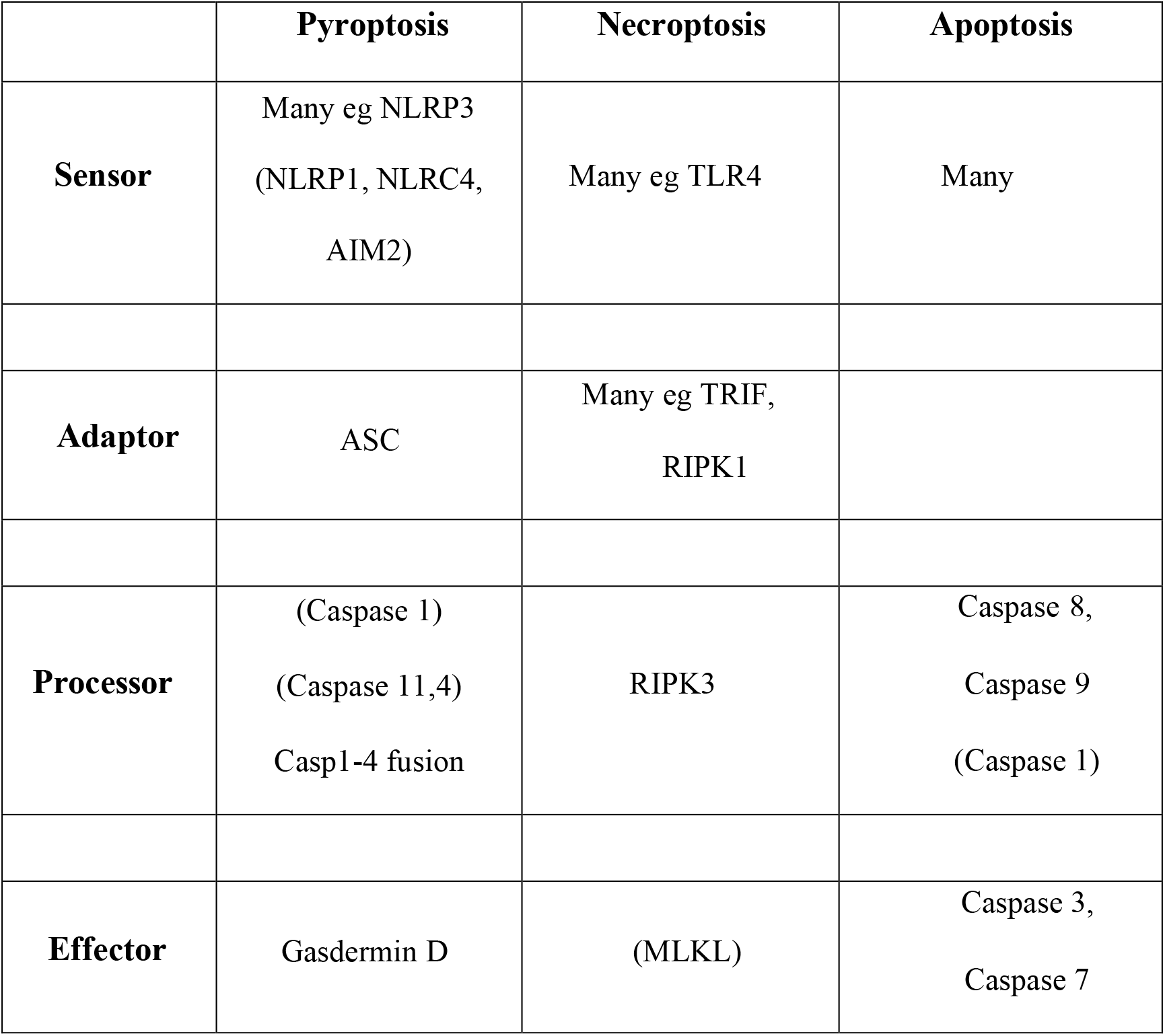
Summary table of cell death pathways in Carnivora.

## Acknowledgments

We would like to thank Vishva Dixit for scientific support, Merone Roose-Girma, Lucinda Tam and Charles Yu, as well as members of the Microinjection, Genetic Analysis, and Animal Resources groups at Genentech for technical assistance in the generation of the DogMo mouse and Scott Snipas for his advice on the recombinant caspase experiments. The immortalized bone marrow derived macrophage lines were a gift from Doug Golenbock and Kate Fitzgerald. Knock-out mouse lines other than DogMo were a gift from Kate Fitzgerald. CEB was supported by a Wellcome Trust Investigator award 108045/Z/15/Z, a Genentech Visiting Professorship, a GSK Immunology Catalyst award and a grant from NodThera. LW is a Wellcome Trust Sir Henry Dale Fellow, ZD was supported by an MRC studentship, JR by a BBSRC studentship. TPM and JPB were supported by a Wellcome Career Development Fellowship awarded to TPM.

## Author contributions

TPM, JPB, LW and JR performed the bioinformatics and evolutionary tracing. ZD, BBV, LJH, performed experiments and analyzed data. SW generated the DogMo mouse, GS, NK, SJW provided critical experimental advice. CEB conceived the study, developed the concept, supervised the research and wrote the manuscript.

## Competing interests

CEB serves on the Scientific Advisory Board of Lightcast and NodThera Inc.

## Data and materials availability

Data in this paper are presented in the main text and supplementary text (raw data can be accessed at https://doi.org/10.17863/CAM.61907).

## STAR Methods

### Data archive

DOI: 10.17632/s8w4ytk5fd.1

https://data.mendeley.com/datasets/s8w4ytk5fd/draft?a=84325eae-bf02-47db-ad78-d1c85b35ba59

### Carnivora gene presence/absence analysis

Genomes of the Carnivora species listed on the Ensembl genome database were examined for the presence of key innate immune system and cell death genes. Genes were first assessed for presence using the standard Ensembl annotation tool. For genes that were absent, we extracted the closest evolutionary relative’s cDNA splice variants (or the human cDNA variants if no relatives could be found in Carnivora) and performed Blastn **(http://www.ensembl.org/Multi/Tools/Blast)** on release 98 of the Ensembl genome database. In some cases, the gene was found but not appropriately annotated. Standard checks were performed to confirm that the gene was a true orthologue of the gene in question. These included 1) producing a bootstrapped neighbour-joining phylogeny and confirming that the gene tree was consistent with the species phylogeny and 2) checking that the gene length was consistent with not containing premature stop codons. The phylogenetic tree was produced in RStudio (v1.2.5001) using the ‘ape’ and ‘seqinr’ packages. The function ‘dist.dna(x, model=”F84”)’ first produced a matrix of pairwise distances from the DNA, ‘nj()’ constructed a tree from this distance matrix and bootstrapping of the tree was carried out using the function ‘boot.phylo()’. A bootstrap value of >80% was taken as support for a particular node. Genes may be missing from genomes due to errors in genome assembly. To verify that genes not found by annotation and Blastn were due to true absence, synteny between the genomes was examined. A queried species genome and the most closely related species genome available containing the investigated gene were compared. The gene coordinates were retrieved from the “Region in detail” tool on Ensembl’s gene location option, this allowed the gene size, location and relation to nearby genes to be obtained. Using this information, synteny maps were generated for species in which the gene is thought to be deleted.

### Cell culture

Murine immortalised bone marrow derived macrophages (iBMDMs) and canine DH82 cells were maintained in Dulbecco’s Modified Eagle Medium (DMEM) containing- 10% FBS, 5 mM L-glutamine, 100 μg/ml streptomycin and 100 units/ml of penicillin at 37°C, 5% CO2. Primary BMDMs were maintained in DMEM containing 10% FCS, 8 mM L-glutamine, 100 μg/ml streptomycin and 100 units/ml penicillin, 20% L929 conditioned medium for 6 days before use. Peripheral blood mononuclear cells (PBMCs) were taken from residual blood samples from dogs admitted to the Queen’s Veterinary School Hospital (Department of Veterinary Medicine, University of Cambridge) and pooled (Table S1). Mononuclear cells were isolated using Histopaque-1077 density gradient medium (Sigma) and SepMate™ mononuclear cell tubes (Stem Cell Technologies). For bacterial infection studies, cell cultures and bacteria were prepared complete medium without antibiotics (DMEM-).

Cells were seeded in 96-well flat-bottomed plates at a concentration of 1×10^6^/ml in DMEM^-^ and incubated overnight at 37°C in 5% CO2. *S*. Typhimurium SL1344 was grown for 18h before sub-culturing to logarithmic growth for 3h as described [1]. Cells were infected with *S*. Typhimurium SL1344 for 1 hour with multiplicity of infection (MOI) 1, 10 or 50. For long time courses (over 2 hours) cell culture medium was replaced with DMEM-cell culture medium supplemented with gentamicin (ThermoFisherScientific) at 50 μg/ml final concentration and further incubated for one hour at 37°C in 5% CO2. For the 6− and 24-hour time-points, 50 μg/ml gentamicin was replaced with fresh DMEM-cell culture medium containing gentamicin at a final concentration of 10 μg/ml. At each time point the supernatant was removed and stored at −80°C until required. In time lapse experiments medium was not replaced to ensure that the field of vision was retained.

### Inflammasome activators and inhibitors

Cells were seeded at 1×10^6^ cells/ml and incubated overnight at 37°C, 5% CO2. Adherent cells were primed with either ultrapure *E.coli* lipopolysaccharide (LPS) (InvivoGen, tlrl-3pelps) at a final concentration of 200 ng/ml for 3 hours or Pam3CSK4 (Pam3) (InvivoGen, tlrl-pms) at a final concentration of 10 μg/ml for 4 hours at 37°C, 5% CO2. Cells were stimulated with nigericin (Sigma-Aldrich, N7143-5MG; 10 μM to 200 μM) for 1 hour. For non-canonical inflammasome activation, cells were incubated with ultrapure LPS at 1 μg/ml or 5 μg/ml final concentration in the presence of FuGENE HD transfection reagent (Promega, E2311) for 16 hours at 37°C, 5% CO2. LPS and transfection reagent were pre-incubated for 15 minutes at room temperature prior to addition to the cells.

### Genome editing

The crRNA designs were performed using Benchling online research platform (https://benchling.com) and crRNA sequences were confirmed using IDT’s Custom Alt-R^®^ CRISPR-Cas9 guide RNA tool (https://eu.idtdna.com/site/order/designtool/index/CRISPR_CUSTOM). Equimolar amounts of tracr- and crRNAs (IDT) were mixed and heated at 95°C for 5 minutes. Ribonucleoprotein (RNP) assembly reaction was performed on duplexes by adding an equimolar concentration of Cas9 endonuclease protein (IDT) to the complex. The RNP complex was then supplemented with equimolar concentration of Alt-R^®^ Cas9 electroporation enhancer (IDT). DH82 cells were resuspended in Nucleofector Solution V containing electroporation supplement (Lonza, VCA- 1003) and 10 μl of RNP was introduced. The nucleofection buffer-cell-RNP mix was then transferred directly into an electroporation cuvette and a single electroporation was performed in an Amaxa Nucleofector 2b (Lonza) using the X-005 program. Cells were rested for 48 hours, after which they were seeded into 96 well plates at single-cell dilution to obtain single cell clones. Monoclones were genotyped as previously described (PMID: 26443227). Briefly, locus-specific primers were designed based on the canine reference genome (CanFam3.1) using Primer-blast and were designed to incorporate 5’ adaptor sequences. First level PCR was performed using genomic DNA extracted from monoclones and the locus-specific primers. Second level MiSeq PCRs were performed using the first round PCR product as template and the universal 96-well Miseq barcode primers, prepared by combining 8 unique forward and 12 unique reverse primers. Second level MiSeq PCR products were pooled, cleaned up and subjected to sequencing using the Illumina MiSeq Reagent Nano Kit v2 300-cycles, #MS-103-1001) according to manufacturer’s instructions on the MiSeq benchtop sequencing systems (Illumina).

### Analysis of sequencing output fastq files

Sequences were analysed using the OutKnocker online analysis tool (http://www.outknocker.org/outknocker2.htm).

### Total protein extraction of cell lysates and cell culture supernatants

Cell pellets were disrupted in ice-cold RIPA solution (150 mM NaCl, 10 mM Tris-HCl, 5 mM EDTA, 1% Triton-X100, 10 mM NaF, 1 mM NaVO4, 20 mM PMSF) supplemented with Protease Inhibitor Cocktail (Sigma-Aldrich) (1:100 dilution) and incubated for 30 minutes on ice. Lysed cells were centrifuged at 14,000 rpm for 15 minutes at 4ºC. Protein concentration was determined using the Pierce™ BCA Protein Assay Kit (ThermoFisherScientific).Standardised amounts of each cell lysate were prepared for immunoblotting by incubating the appropriate volume of sample with Pierce™ Lane Marker (5x) Reducing Sample Buffer (Thermo Fisher) for 10 minutes at 100°C. After denaturation lysates were cooled on ice followed by brief centrifugation and subjected to SDS-PAGE.

### Total protein extraction from cell culture supernatants

Total protein was extracted from clarified supernatants using methanol/chloroform extraction method. Samples were then centrifuged at 16,000 × g for 12 minutes at 4°C. The intermediate phase containing the precipitated protein was carefully aspirated and centrifuged for 10 minutes at 4°C. Protein pellets were washed twice in ice-cold methanol. In between washes the resuspended pellets were centrifuged at 16,000 × g for 5 minutes at 4°C. After the final wash, the methanol was carefully removed, and the pellets were allowed to dry. Pellets were re-suspended in Laemmli 2 × concentrate sample buffer (Sigma Aldrich), heated at 100°C for 10 minutes and stored at −80°C until required.

### Western blotting

Denatured proteins were separated by SDS-PAGE using 12% tris-glycine polyacrylamide gels and subsequently transferred to nitrocellulose membranes followed by incubation with primary (anti-ASC #AG-25B-006-C100, anti-canine IL-1β AF3747, anti-caspase-1 p10 sc-514, anti-caspase-11 sc-56038 or anti-GSDMD ab209845) and HRP-conjugated secondary antibodies. Proteins were detected using Western Lightning Plus-ECL Substrate (Perkin Elmer, NEL103E001EA) and protein bands were visualised using a GeneGnome chemiluminescence imager (Syngene, GeneGnome XRQ).

### LDH release quantification

Supernatants were assayed for LDH release at indicated time points using the CytoTox 96^®^ Non-Radioactive Cytotoxicity Assay kit (Promega, G1780) per the manufacturer’s instruction. Absorbance values (absorbance 490 nm and 680 nm) were measured on either a PHERAstar or CLARIOstar Microplate Reader (both BMG Labtech). Percentage cell death values are expressed relative to total LDH release induced by detergent lysis.

### Propidium iodide uptake

DH82 were seeded into 96-well black/clear bottom microplates at 5 × 10^4^ cells/well in complete DMEM and incubated overnight. Cells were either unprimed or primed with 200 ng/ml LPS for 3 hours prior to stimulation with 100 μM nigericin. Cells were infected with *Salmonella* at the indicated MOIs in imaging medium (Opti-MEM containing GlutaMax, HEPES and 10% FBS). Propidium iodide (PI) was used at a final concentration of 2 μg/ml. Triton X-100 (0.2% w/v) in imaging medium was added to control wells. Measurements were taken every 10 minutes on a CLARIOstar microplate reader set to 37 °C. After the first kinetic window (1 hour), the plate was removed and 100 μM nigericin added to the LPS-primed wells, or in the *Salmonella*-infected wells, the medium was replaced with imaging medium supplemented with 50 μg/ml gentamicin, and fluorescence readings recorded every 10 minutes for a further 23 hours. The percentage PI uptake was normalised against the unstimulated fluorescence recordings at the start of each kinetic window (0%) and the maximal PI uptake (100 %) induced by Triton X-100 (0.2 %).

### Cytokine measurement using ELISA

Mouse IL-1β and canine IL-1β released into cell culture supernatant was measured using Mouse IL-1β ELIOptEIA™ (BD Biosciences, 559603) and Canine IL-1β/IL-1F2 DuoSet (R&D Systems, DY3747) respectively, as described in the manufacturer’s instructions. Absorbance was read at 450 nm wavelength (570 nm correction) using either a PHERAstar or CLARIOstar Microplate Reader (BMG, Labtech).

### Sample preparation for immunofluorescent staining and imaging

Murine and canine iBMDMs and primary macrophages were seeded at a density of 2×10^5^ cells/well on an 8-well chamber slide (ThermoFisher) and incubated overnight at 37°C, 5% CO2. Following ligand stimulation, cells were incubated with either FAM FLICA caspase-1, caspase-8or poly-caspase (all Bio-Rad) according to the manufacturer’s instructions. Post stimulation, cells were fixed in 4% paraformaldehyde in PBS (Thermo Fisher Scientific, 28908) and incubated with anti-ASC (AL177) pAb (Adipogen, AG-25B-0006-C100) followed by incubation with goat-anti-rabbit-IgG-AlexaFluor488 (Invitrogen, A11034) or -AlexaFluor568 (Invitrogen, A11036) secondary antibodies. Cells were counterstained with nuclear Hoechst 33342 (Invitrogen, H3570) labelling solution followed by mounting in VECTASHIELD Antifade Mounting Medium (H-1000, Vector) and imaged using an Inverted Fluorescence Microscope (Leica DM IRM).

### Recombinant protein expression and *in vitro* caspase activity assays

Constructs encoding mouse pro-IL-1β, as well as CARD deleted modified versions of caspase-1 and caspase-11 in pET29b(+) containing a C-terminal 6×His tag were previously described[2]. The DNA encoding the CARD deleted version of dog caspase−1/−4 containing a C-terminal 6×His tag was purchased from Integrated DNA technologies (IDT, San Diego, CA) and cloned into pET29b(+) by using Nde I and Xho I. Gasdermin D sequence was amplified from a pET29b(+) construct using primers to add an N-term 8xHis and cloned into pET15b using Nde I and Xho I restriction enzymes. Proteins were expressed in BL21(DE3) *E. coli* cultures and purified as previously described (Ramirez et al., 2018). Protein concentration of pro-IL-1β and gasdermin D was calculated by absorbance at 280 nm and the concentration of caspases was calculated by active site titration with z-VAD-fmk. Activity assays were performed in 100 μl final volume and assay buffer consisted of 20 mM PIPES, 10% sucrose, 100 mM NaCl, 0.1% CHAPS, 1 mM EDTA, 10 mM DTT and 0.75 M sodium citrate. The fluorogenic peptide substrates Ac-Trp-Thr(Bzl)-His-Asp-ACC and Ac-Tle-Bpa-His(Bzl)-Asp-ACC and Ac-WEHD-AFC was purchased from BACHEM. For *k*cat/KM determination, the concentration of fluorogenic peptide substrates varied in the range of 5-200 μM. Reactions were monitored for 30 minutes at 37°C in a CLARIOstar plate reader (BMG LabTech). The ACC fluorophore was detected at excitation/emission 355/460 nm and AFC at 400/505 nm. Reaction velocity was calculated using MARS data analysis software (BMG LabTech) and the kinetic parameters with Prism 7 (GraphPad) using the Michaelis-Menten equation. Recombinant inflammatory caspases were subjected to 2-fold dilution series and incubated for 30 minutes at 37°C with 4 μM gasdermin D or pro-IL1β. Reactions in a 60 μl final volume were performed using assay buffer minus sodium citrate. After incubation, reactions were terminated by the addition of 30 μl of 3 × SDS loading buffer and heated at 95°C for 5 minutes. Reaction products were separated on 4-12% Bis-Tris polyacrylamide gels and stained with Instant Blue (Expedeon).

### Generation of a mouse with a*Casp1*-*Casp4* gene fusion

To generate a mouse allele corresponding to the dog caspase gene fusion, a CRISPR strategy with two sgRNAs was used to generate a 20,524 bp deletion (GRCm38/mm10 chr9:5,302,869-5,323,392). The 5’ sgRNA (binding to chr9:5,302,865-5,302,884 reverse strand) is located in *Casp1* intron 4 and the 3’ sgRNA (binding to chr9:5,323,374-5,323,393) is located in *Casp4* intron 3. The deletion thus created an in-frame fusion between *Casp1* exons 1-4 and *Casp4* exons 4-9, resulting in a fusion protein similar to the dog fusion protein (see protein alignment). Cas9 mRNA and synthetic sgRNAs (Synthego) were co-microinjected into C57BL/6N mouse zygotes and resulting mosaic founders positive for the large deletion were analysed for absence of off-targets essentially as described and subsequently bred to C57BL/6N for transmission of the fusion allele. Homozygous mice were used in this study (Anderson et al., 2018; Sievers et al., 2011).

## Supplementary Materials

**Figure S1:**
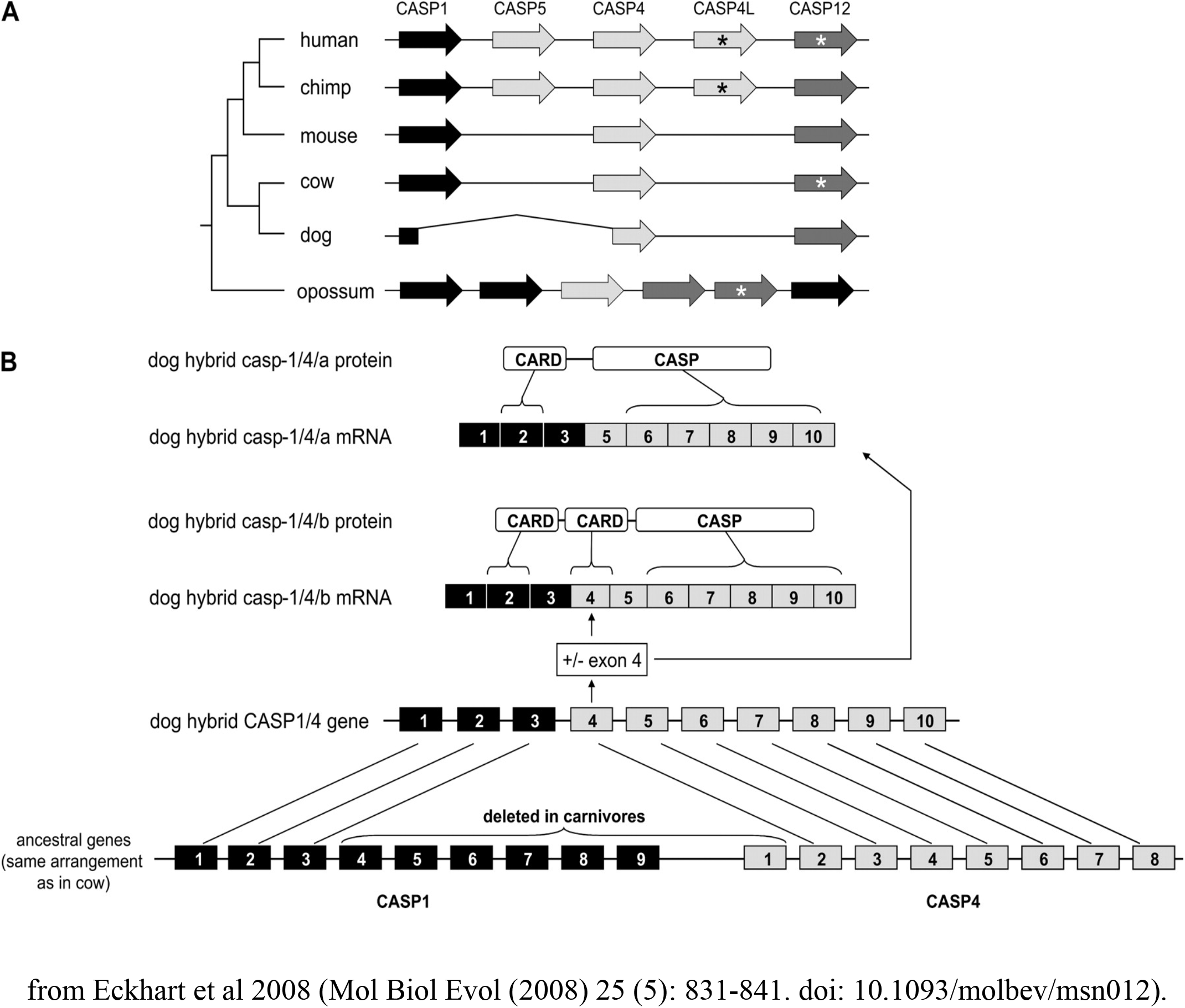
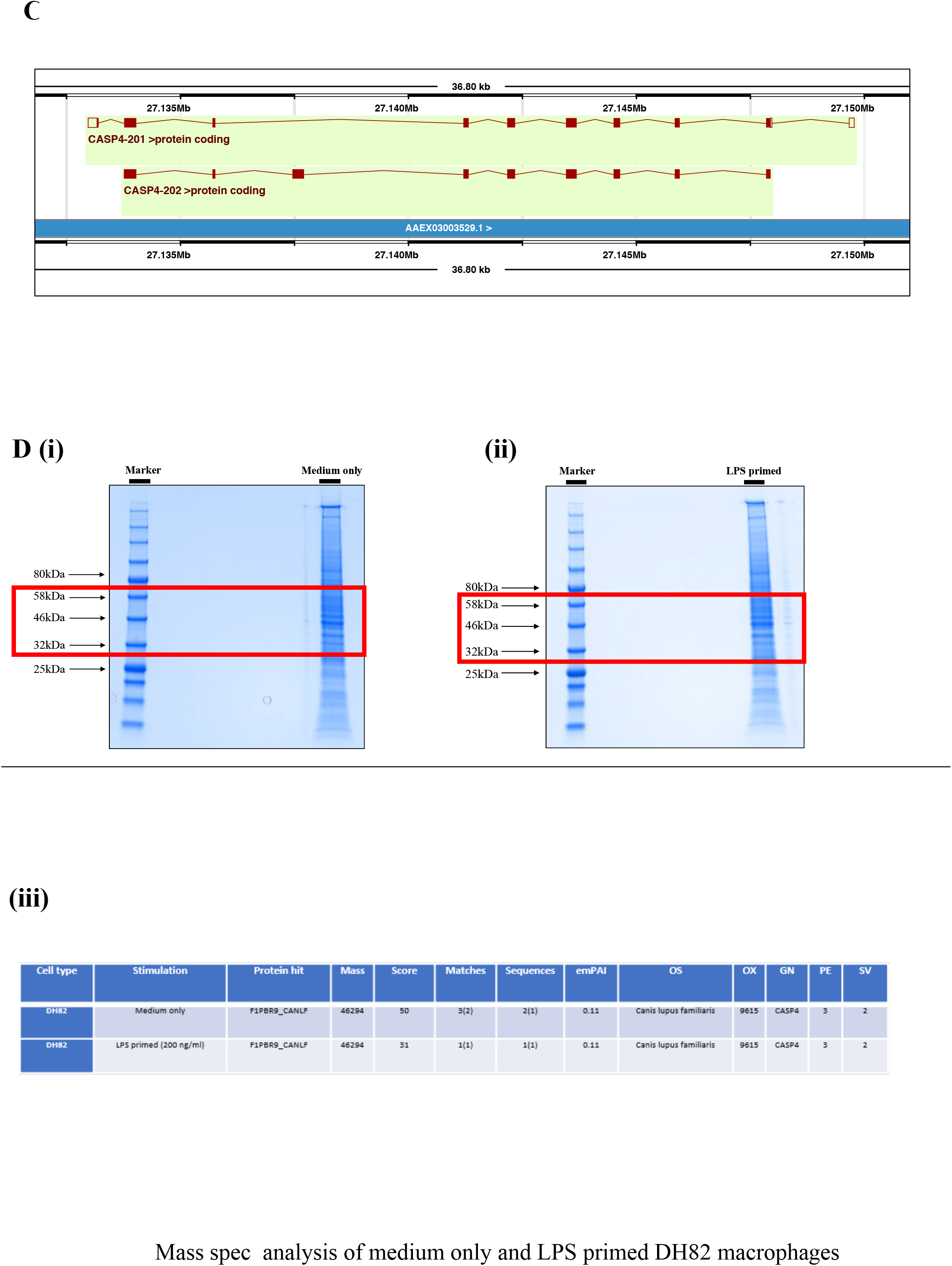
The dog caspase−1/−4 hybrid (A, B) Domain organisation of the dog caspase−1/4/11 hybrid protein. The gene encoding the canine hybrid protein can produce two alternative transcripts. (Transcript A) The larger transcript contains two tandem CARDs and a caspase domain. The first CARD domain shows high sequence homology to the CARD domain of human and murine caspase-1. While the second CARD domain shows high sequence similarities to the CARD domain of human caspase−4/−5 and murine caspase-11. The caspase domain shows high similarities to the caspase domain of human caspase−4/−5 and murine caspase-11. (Transcript B) By contrast, the shorter transcript excludes the second CARD domain, resulting in a gene sequence consisting of a CARD domain similar to human and murine caspase-1 and the caspase domain similar to human caspase−4/−5 and murine caspase-11. **(C)** Ensembl gene build annotation of the hybrid gene (ENSCAFG00000014860) shows the predicted protein coding exons. The identified catalytic residue is present in both transcripts in exon 6. Mass spectrometry analysis of full protein extracts from **(D(i))** non-stimulated (medium only) and **(ii)** LPS primed DH82 cell lysates were cut out between the 32 kDa and 58 kDa molecular weight markers (area marked with red rectangles) and prepared for mass spectrometry analysis by the Cambridge Centre for Proteomics (CCP) Core Facility. **(iii)** Mass spectrometry analysis showed constitutive expression of the hybrid gene.

**S2.**
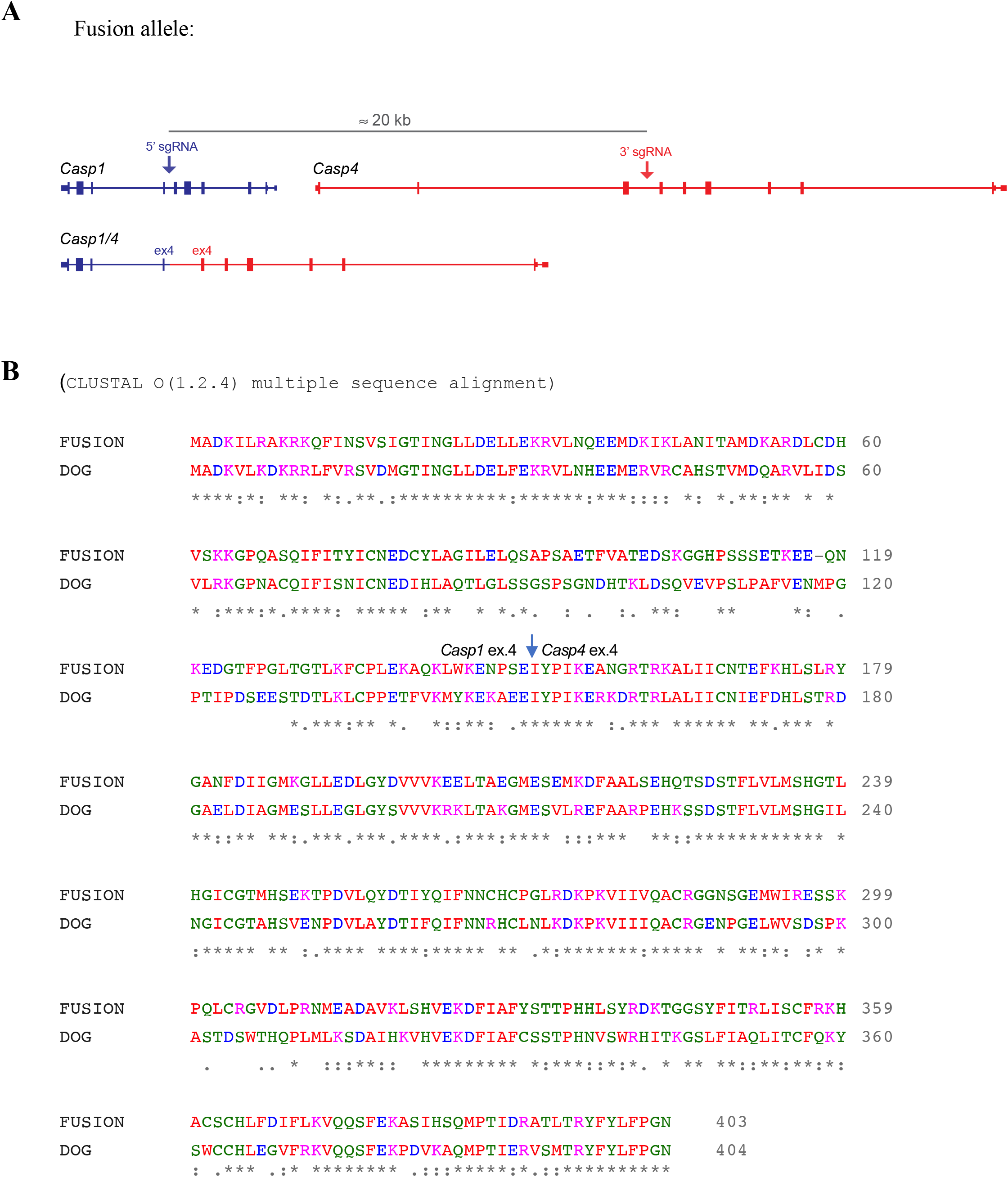
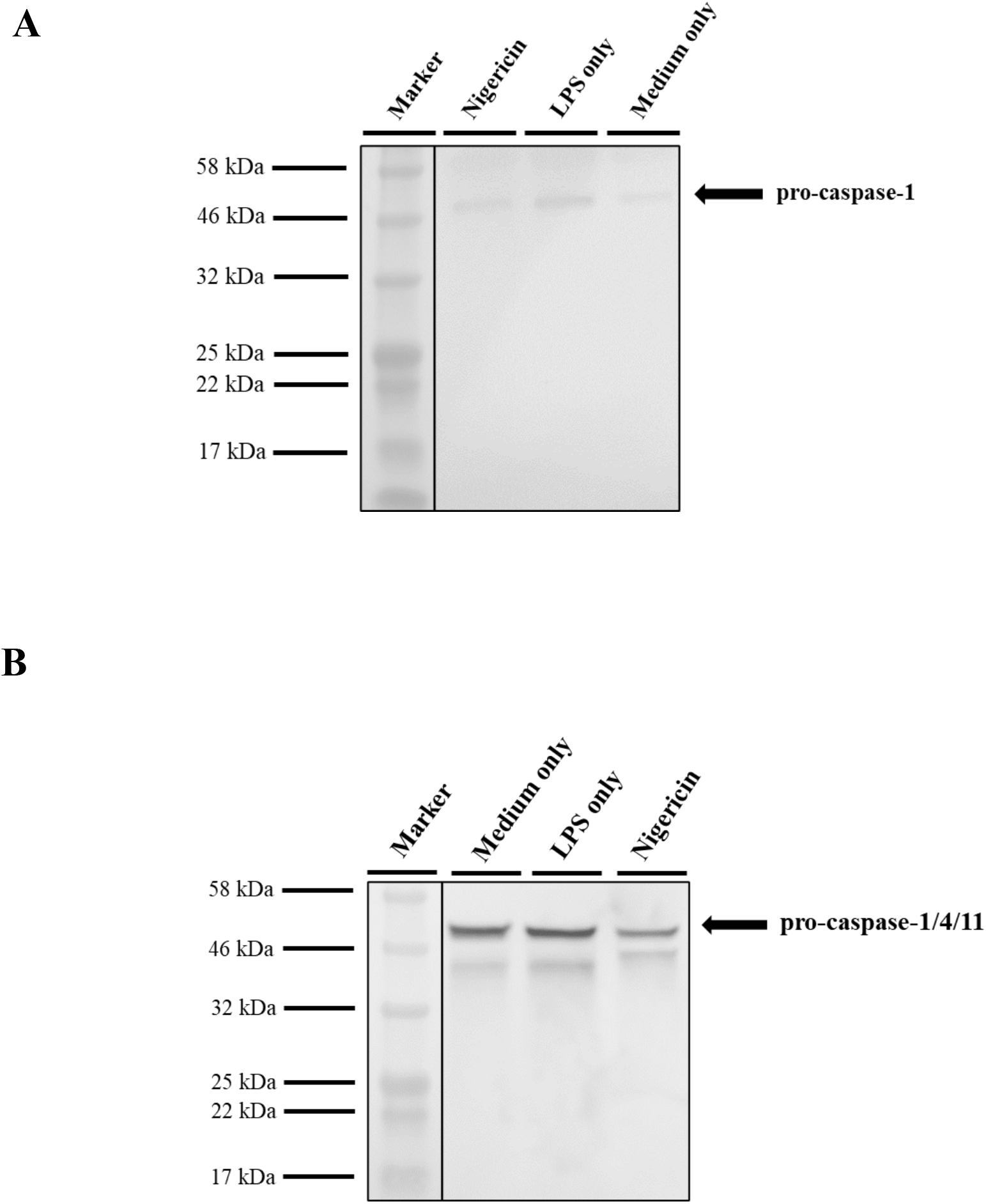
Strategy for the generation of the DogMo mouse. **(A)** To generate a mouse allele corresponding to the dog caspase gene fusion, a CRISPR strategy with two sgRNAs was used to generate a 20,524 bp deletion (GRCm38/mm10 chr9:5,302,869-5,323,392). The 5’ sgRNA (binding to chr9:5,302,865-5,302,884 reverse strand) is located in Casp1 intron 4 and the 3’ sgRNA (binding to chr9:5,323,374-5,323,393) is located in Casp4 intron 3. **(B)** The deletion thus created an in-frame fusion between Casp1 exons 1-4 and Casp4 exons 4-9, resulting in a fusion protein similar to the dog fusion protein. **Immunoblotting showing the expression of the caspase−1/−4 hybrid gene in DogMo cells**. Murine primary WT and DogMo cells were primed with LPS (200 ng/ml for 3 hours) followed by incubation with nigericin (20 μM for 1 hour). Caspase-1 and the hybrid caspase−1/−4 protein expression was confirmed by western blotting of cell lysates using murine specific caspase-1 and caspase-11 antibody. respectively. Representative immunoblots are from a single experiment

**S3.**
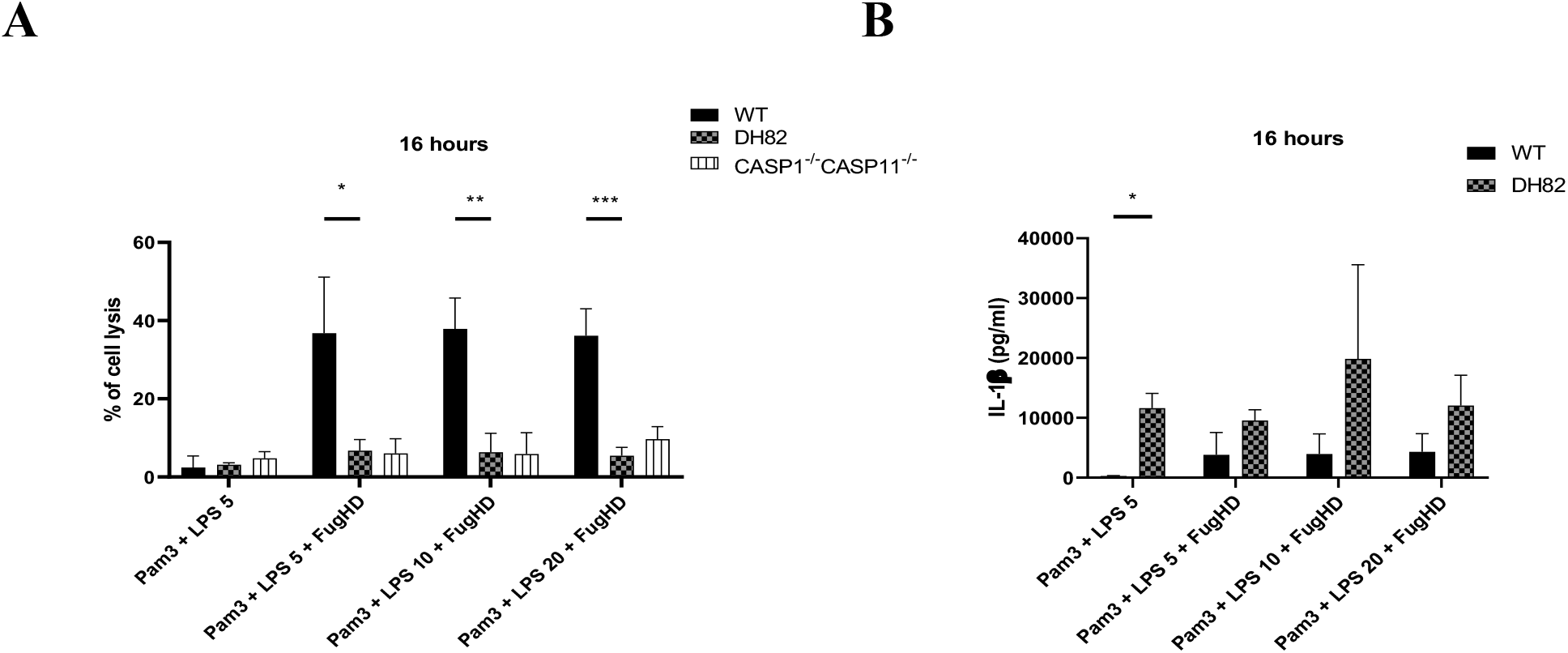
Comparison of cell death and IL-1β production from canine DH82, WT and CASP1/CASP11 double knock out mice in response to cytosolic LPS transfection. Immortalised murine WT, CASP1−/−CASP11−/− and canine DH82 cells were primed with Pam3CSK4 (10 μg/ml for 4 hours) then stimulated with either LPS (5 μg/ml) alone or in conjunction with FuGENEâHD transfection reagent. Percentage of cell lysis induced was determined by measuring LDH release into the supernatant, while IL-1β release was measured using ELISA method. Data shown is pooled from three independent experiments. Error bars represent the standard error of means (SEM) of triplicate wells.

**S4.**
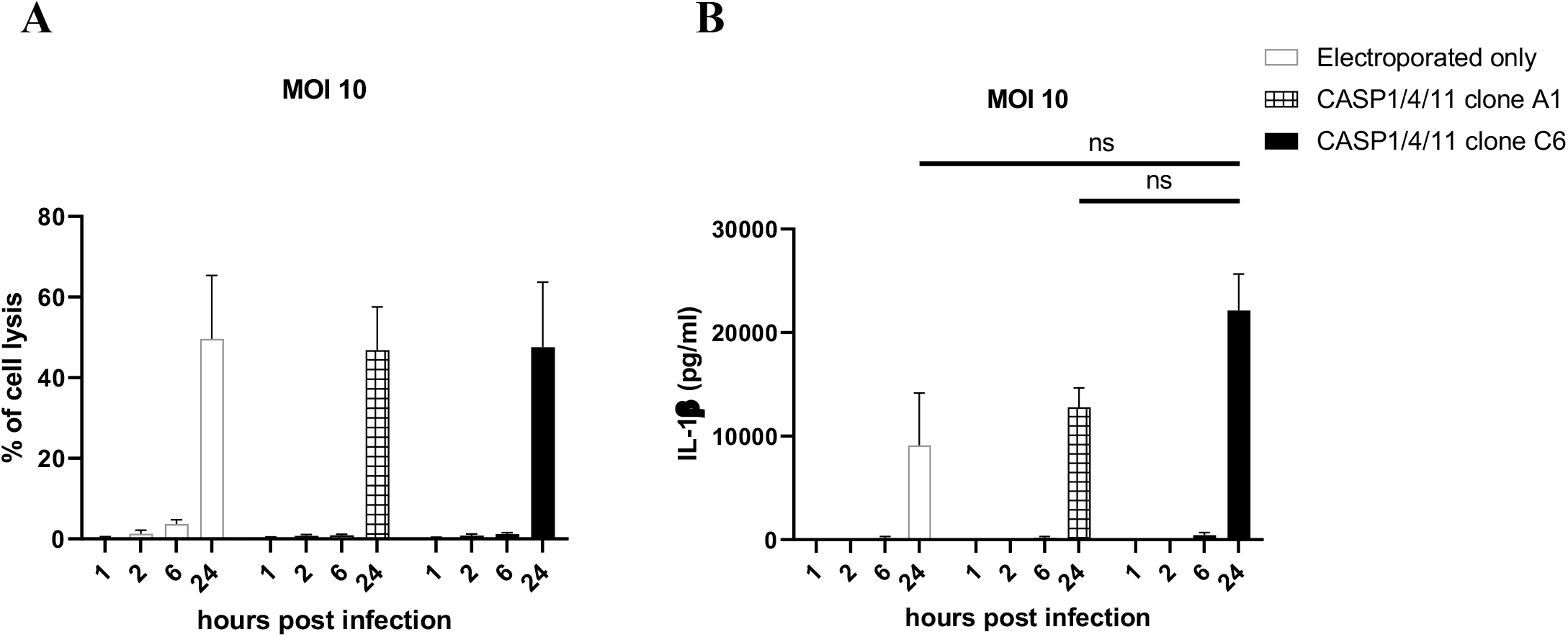
SCRISPR/Cas9 deletion of the caspase−1/−4 from DH82 does not affect IL-1β production in response to inflammasome stimulation by *S*Tm infection. Caspase−1/−4 knock-out clones A1, C6, non-electroporated and electroporated wild-type DH82 cells were infected with *S.* Typhimurium at MOI of 10. **(A)** Percentage of cell lysis induced was determined by measuring the relative amount of LDH release into the supernatant, while **(B)** IL-1β release was measured using ELISA method. Data shown is pooled from three independent experiments. Error bars represent the standard error of means (SEM) of triplicate wells.

**Table S1.**

| Identifier | Breed | Age | Reason for blood test |
| --- | --- | --- | --- |
| 1379 2901 | Whippet | 9 years | Chemo monitoring |
| 1379 3501 | Retriever | 12 years | Monitoring of treatment for anal sac adenocarcinoma |
| 1379 4801 | Cocker Spaniel | Unknown | Monitoring of immune-mediated haemolytic anaemia |
| 1379 5301 | Staffordshire Bull Terrier | 11 years | Chemo monitoring |
| 1379 5401 | Labradoodle | 12 years | Screening |
| 1379 6201 | Greyhound | 6 years | Chemo monitoring |

B
| Identifier | Breed | Age | Reason for blood test |
| --- | --- | --- | --- |
| 1386 0901 | Pug | 5 years | Protein-losing enteropathy |
| 1386 1301 | Cacapoo | 4 years | Immune Mediated Thrombocytopenia re-check |
| 1386 3601 | Staffordshire Bull Terrier | 8 years | Screening |

## Notes

https://www.example.com

### Competing Interest Statement

CEB served as a member of the GSK Immunology Catalyst and currently serves on the Scientific Advisory Board of Lightcast and NodThera Inc.

